# PI3K/AKT signalling orchestrates ICM maturation and proper epiblast and primitive endoderm specification

**DOI:** 10.1101/2023.06.21.545867

**Authors:** Anna Geiselmann, Adèle Micouin, Sandrine Vandormael-Pournin, Vincent Laville, Sébastien Mella, Pablo Navarro, Michel Cohen-Tannoudji

**Affiliations:** Institut Pasteur, Université Paris Cité, CNRS UMR3738, Epigenomics, Proliferation, and the Identity of Cells, Department of Developmental and Stem Cell Biology, F-75015 Paris, France; Sorbonne Université, Complexité du Vivant, F-75005, Paris, France; Institut Pasteur, Université Paris Cité, CNRS UMR3738, Early Mammalian Development and Stem Cell Biology, F-75015, Paris, France; Université Paris Cité, BioSPC, F-75013, Paris, France; Department of Developmental and Stem Cell Biology, Institut Pasteur, CNRS UMR 3738, F-75015, Paris, France; Institut Pasteur, Bioinformatics and Biostatistics Hub (C3BI), F-75015, Paris, France

**Author notes:** Corresponding author and lead contact: Michel Cohen-Tannoudji, Early Mammalian Development and Stem Cell Biology, Institut Pasteur, CNRS UMR 3738, 25 rue du Dr. Roux, F-75015, Paris, France; website: https://research.pasteur.fr/en/team/group-michel-cohen-tannoudji.

**Keywords:** Mouse preimplantation embryo, Epiblast, Primitive endoderm, Inner cell mass, Lineage specification, NANOG, GSK3, FOXO3, mTOR

## Abstract

The inner cell mass (ICM) of early mouse embryos is specified into Epiblast (Epi) and primitive endoderm (PrE) lineages during blastocyst formation. The antagonistic transcription factors (TFs) NANOG and GATA6 in combination with FGF/ERK signalling are central actors in ICM fate choice. However, what initiates the specification of ICM progenitors and whether other factors are involved in this process is not fully understood yet. Here, we show that PI3K/AKT is constitutively active during preimplantation development. Using pharmacological inhibition, we demonstrate that PI3K/AKT enables the formation of a functional ICM capable of giving rise to both the EPI and the PrE: it maintains the expression of the TF NANOG, which specifies the EPI, and confers responsiveness to FGF4, which is essential for PrE specification. Our observations thus identify PI3K/AKT signalling as an upstream regulator orchestrating the molecular events required for both EPI and PrE specification.

## Introduction

During mouse preimplantation development, transcription factors (TFs)-controlled networks and coordinated cell signalling pathways orchestrate the formation of the first three embryonic lineages composing the implanting blastocyst. The pluripotent epiblast (Epi) will give rise to the foetus and the extraembryonic mesoderm. The trophectoderm (TE) and primitive endoderm (PrE) will contribute to the placenta and the endoderm layers of the yolk sacs, respectively. TE and inner cell mass (ICM) fates are established during the 4^th^ and 5^th^ division. Shortly after, during blastocyst formation, individual ICM cells acquire PrE or Epi fate in an asynchronous and mutually exclusive manner but without predefined spatial patterns leading to mutually exclusive expression of lineage specific markers in a ‘salt-and-pepper’ distribution (Chazaud et al., 2006; Plusa et al., 2008). Before segregating the expression of lineage-specific TFs, ICM progenitors coexpress Epi-(NANOG and SOX2) and PrE-specific (GATA6) TFs. While a subset of the cells will maintain NANOG and downregulate GATA6, specifying the Epi, others will silence NANOG and then other pluripotency factors (e.g. SOX2 and OCT4), activate PrE-specific TFs including SOX17, GATA4 and SOX7 (Artus et al., 2011), and specify the PrE. In this setting, NANOG and GATA6 act as major regulators of their respective lineage (Bessonnard et al., 2014; Frankenberg et al., 2011; Schrode et al., 2014), with the Fibroblast Growth Factor (FGF)/Extracellular signal-Regulated Kinase (ERK) signalling pathway also playing a major role in driving PrE fate acquisition. Indeed, inactivation of genes of the pathway or treatment with inhibitors prevents PrE formation (Chazaud et al., 2006; Kang et al., 2013; Kang et al., 2017; Molotkov et al., 2017; Nichols et al., 2009; Yamanaka et al., 2010), whereas overactivation of the pathway can bias the entire ICM towards the PrE lineage (Saiz et al., 2016; Yamanaka et al., 2010). In fact, Epi progenitors are established first (Bessonnard et al., 2014; Bessonnard et al., 2017; Saiz et al., 2016) and the upregulation of *Fgf4* expression triggers PrE fate in surrounding uncommitted ICM cells by inducing ERK signalling (Azami et al., 2019; Frankenberg et al., 2011; Kang et al., 2017). In *Gata6* mutant embryos, NANOG is expressed in all ICM cells, and no PrE progenitors are formed, even after the addition of exogenous FGF4, which fails to rescue expression of SOX17 and GATA4 (Bessonnard et al., 2014; Schrode et al., 2014). Thus, GATA6 and FGF/ERK signaling are acting in synergy to promote PrE identity.

Overall, relatively simple Gene Regulatory Networks, centered around direct mutual inhibition between NANOG and GATA6 and integrating positive and negative feedback from the FGF/ERK pathway, allow to model many aspects of Epi and PrE cell fate decision (Mot et al., 2016; Saiz et al., 2019; Tosenberger et al., 2017). However, how the initial separation between Epi and PrE fate is controlled remains incompletely understood. Our previous results showed that NANOG is required to initiate Epi cell specification (Allègre et al., 2022). Regulation of NANOG levels in ICM progenitors is therefore central to drive the initial bias towards Epi or PrE fate. Noteworthy, most of our current knowledge on NANOG regulation comes from work on pluripotent stem cells and much less is known about transcriptional and post-transcriptional regulation of *Nanog in vivo* in Epi cells, especially in ICM progenitors which express both GATA6 and NANOG and are endowed with different properties and potentialities.

The Phosphoinositide 3-kinase (PI3K)/ protein kinase B (AKT) signalling pathway plays critical roles in metabolism, proliferation and survival of most cell types. Besides these essential cellular functions, PI3K/AKT is key for the maintenance of pluripotency in mouse and human pluripotent stem cells (Yu and Cui, 2016). Acting downstream of LIF and IGF1, PI3K/AKT activity is required to sustain expression of pluripotency factors including TBX3 and NANOG (Hishida et al., 2015; Niwa et al., 2009; Storm et al., 2007; Storm et al., 2009; Wamaitha et al., 2020). There is evidence that the pathway is active in preimplantation embryos (Halet et al., 2008; Riley et al., 2005), where it is likely to mediate the actions of trophic factors present in maternal fluids or produced by the embryos themselves. Indeed, treatments with IGF-1/Insulin, or pharmacological inhibitors, point to the involvment of the pathway in growth and survival of preimplantation embryos in various mammals, including humans (Campbell et al., 2012; Lin et al., 2003; Lu et al., 2004; Ramos-Ibeas et al., 2019; Wamaitha et al., 2020). We further showed that, in the mouse embryo, PI3K/AKT/mTOR is critically required for the survival of both Epi and PrE after their specification (Bessonnard et al., 2019). How PI3K/AKT activity is regulated in the embryo and whether it contributes to the specification and establishment of Epi or PrE fates remains largely unknown.

In this study, we addressed the role of PI3K/AKT signalling during the time-window of Epi and PrE specification in the mouse embryo. Using phosphorylated ribosomal protein S6 (pRPS6) as a readout of the pathway, we demonstrated that PI3K is active during all preimplantation development and increases in mid/late blastocysts. By pharmacologically modulating its activity, we showed that PI3K regulates the levels of NANOG and other pluripotency TFs in ICM progenitors and confers the competence to respond to FGF4.

## Results

### PI3K/AKT pathway is active during preimplantation development

To monitor PI3K/AKT activity during preimplantation development, we analyzed the levels of phosphorylated ribosome protein S6 (pRPS6), a well-established downstream target and widely used readout of PI3K/AKT pathway activity (Meyuhas, 2015). We first quantified the levels of total and phosphorylated RPS6 from the two-cell-stage to the late blastocyst stage (embryonic day (E) 4.0; Figure 1A) embryos. As expected for an integral component of the small ribosomal subunit, RPS6 was detected in the cytoplasm of all blastomeres and at all stages. During preimplantation development, total RPS6 levels increased and peaked at early blastocyst (E3.25) stage (Figure 1A and S1A). Similar to total RPS6, pRPS6 was detected in all cells and at all stages and up to E3.25 followed the same trend than total RPS6 expression (Figure 1A and S1A). Accordingly, the ratio of pRPS6 over total RPS6 cytoplasmic levels was quite homogeneous between blastomeres and remained stable over this developmental period (Figure 1B). In contrast, heterogeneous pRPS6 levels were observed at later stages in both inner and outer cells (Figure 1A-B). To address whether variation in pRPS6 levels in inner cells was correlated with Epi and PrE fates, we quantified pRPS6, NANOG and GATA6 levels in inner cells of E3.25, E3.75 and E4.0 embryos. We found no correlations between RPS6 phosphorylation and NANOG or GATA6 levels (Figure S1B) suggesting that fixed levels of pRPS6 are not linked to ICM, Epi or PrE identities. Heterogeneous pRPS6 levels may rather reflect various cellular processes including the PI3K/AKT-mediated survival of Epi and PrE progenitors once these are specified (Bessonnard et al., 2019).

**Figure 1:**
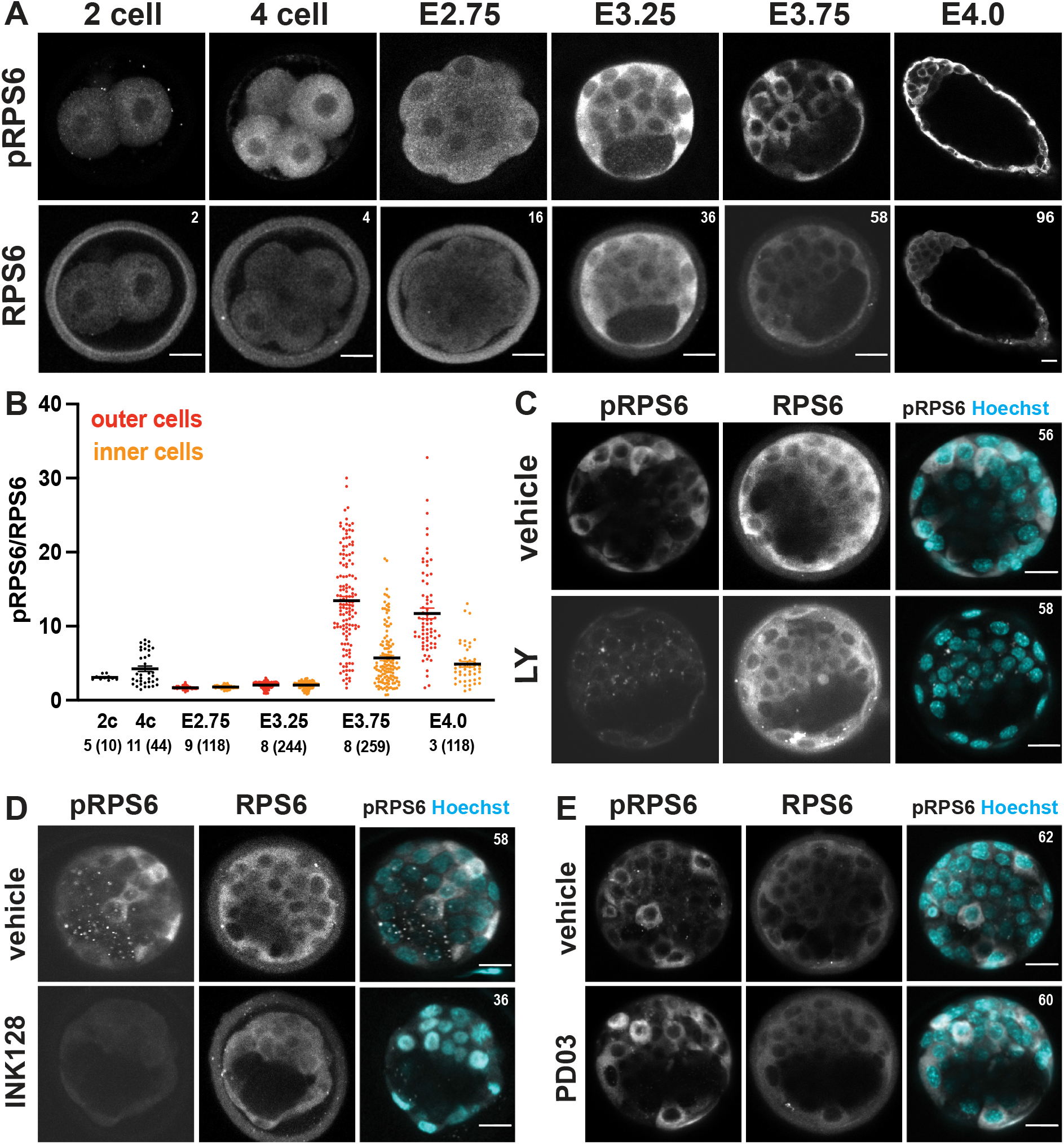
pRPS6 staining reveals two phases of PI3K/AKT activity during preimplantation development. (A) Immunostaining of pRPS6 and RPS6 (grey) from 2 cell stage to E4.0. Distinct pRPS6 levels are visible in E3.75 and E4.0. (B) Ratio of pRPS6/RPS6 quantification levels in inner and outer cells from 2 cell stage to E4.0. Number of embryos and analyzed cells per stage are indicated at the bottom. (C-E) Immunodetection of pRPS6 and RPS6 (grey) in embryos cultured from E2.75 for 24h in (C) the PI3K inhibitor, (D) the mTORC1 and mTORC2 dual inhibitor INK128 or (E) the ERK inhibitor PD03. Data are from one experiment, representative of two (A-B, D-E) and four (C) independent experiments. Pictures correspond to a single (A) or the projection of 5 (D-E) optical slices. Number of cells per embryo are indicated on the top right of images. Scale bar: 20µm.

RPS6 phosphorylation is mainly catalyzed by S6 kinase (S6K) upon mammalian target of rapamycin complex 1 (mTORC1) activation. However, it can also occur through other pathways including 90-kDa rpS6 Kinase (RSK) upon MEK/ERK activation. To determine which pathways are responsible for RPS6 phosphorylation during preimplantation development, we cultured E2.75 embryos for 24h with various inhibitors: the PI3K inhibitor LY294002 (LY), the mTOR inhibitors Rapamycin and INK128, or the Mitogen-activated protein kinase (MEK) inhibitor PD0325901 (PD03). We observed a complete disappearance of pRPS6 staining upon PI3K or mTOR but not MEK inhibition (Figure 1C-E, S1C). Collectively, our data show that PI3K/AKT/mTOR signalling is constitutively active throughout preimplantation development and becomes more intense and heterogeneous once the first three embryonic cell lineages are established.

### PI3K/AKT maintains NANOG in ICM progenitors

Next, we monitored the consequences of PI3K inhibition on NANOG expression by treating embryos with LY during various time windows. NANOG expression is first detected at the 8-cell stage (Plusa et al., 2008). To test a possible role of PI3K/AKT at the onset of NANOG expression, we treated 2-cell stage embryos for 24h until they reached the 8-cell stage. Treated embryos developed normally and showed NANOG staining (Figure S2A) indicating that PI3K/AKT activity is not required for the initiation of *Nanog* expression. We then treated E2.75 morulae for 24h, a period during which ICM progenitors are produced and initiate their specification into Epi and PrE. Strikingly, we observed that although both LY and vehicle treated embryos progressed similarly to mid-blastocyst stage (∼50-64 cells), a dramatic reduction in the number of NANOG-positive was observed upon PI3K inhibition (Figure 2A). Quantification of NANOG and GATA6 levels in inner cells showed a strong reduction of NANOG levels upon PI3K inhibition while GATA6 levels were hardly affected (Figure 2B). Treatment of E2.75 morulae for a shorter period of 16 hours, until they reached early-blastocyst (∼30-38 cells) stage, or of E3.0 early blastocysts (∼27-32 cells) for 8 hours gave similar results (Figure S2B-E). In contrast, E3.25 embryos (∼47-55 cells) treated for 8 hours showed no strong alteration of NANOG levels (Figure 2C-D). Together, these observations suggest that PI3K/AKT inhibition impacts NANOG levels during the birth of ICM progenitors and the initial phase of Epi/PrE specification, but not at a later phase.

**Figure 2:**
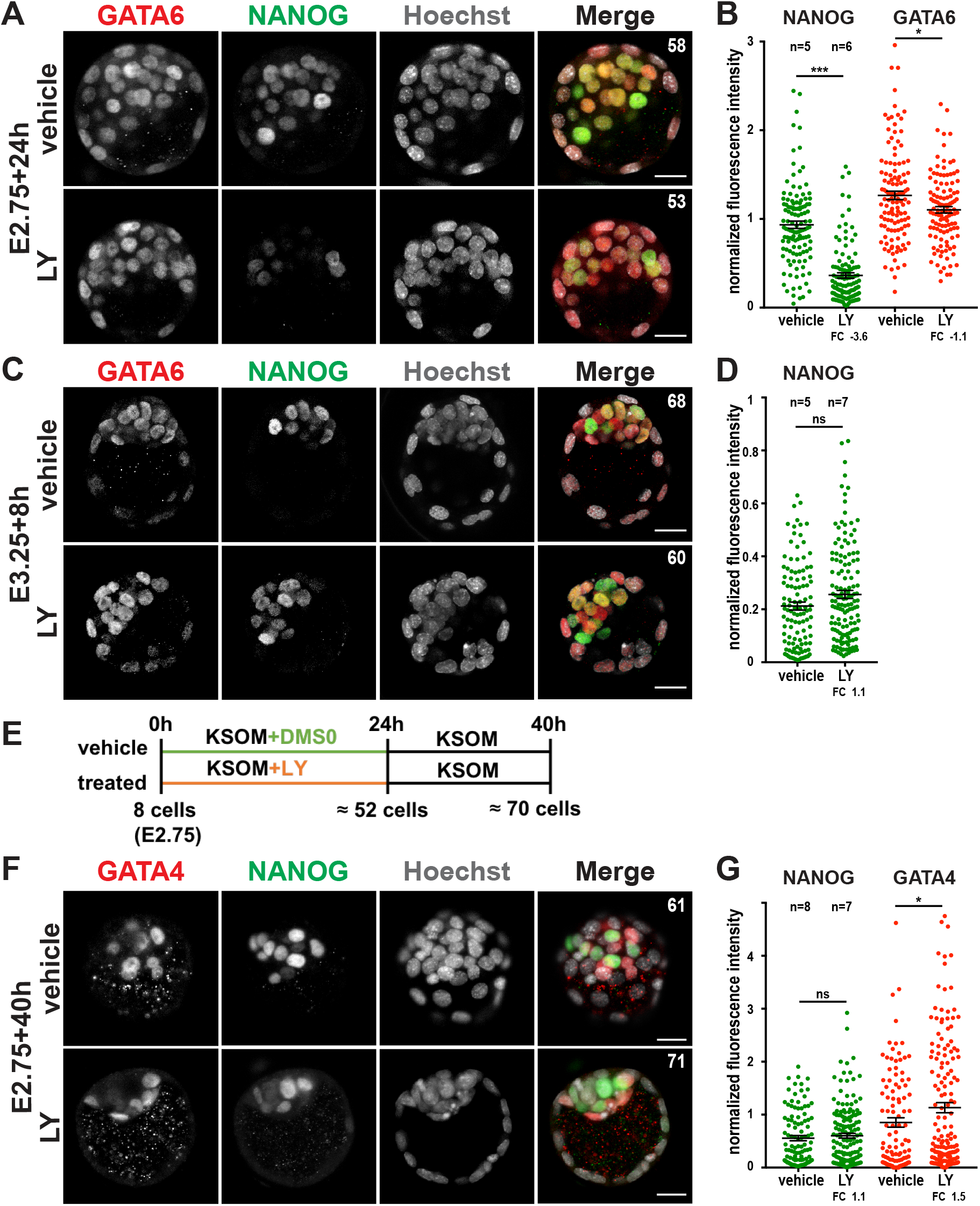
PI3K inhibition during ICM specification leads to NANOG reversible downregulation. (A) Immunodetection of GATA6 (red) and NANOG (green) and (B) quantification of GATA6 (red) and NANOG (green) in embryos cultured for 24h from E2.75. (C) Immunodetection of GATA6 (red) and NANOG (green) and (D) quantification of NANOG (green) in embryos cultured for 8h from E3.25. (E) Schematic of the release experiment shown in (F-G). E2.75 embryos were cultured for 24h starting in KSOM containing either DMSO (vehicle) or LY (treated) before being transferred to KSOM alone for an additional 16h before fixation. (F) Immunodetection of GATA4 (red) and NANOG (green) and (G) quantification of GATA4 (red) and NANOG (green) in the embryos at the end of the release experiment. Graph shows results of two independent experiments. Pictures correspond to the projection of 5 optical slices. Data are from one experiment, representative of >10 (A-B) and 3 (C-G) independent experiments. Number of cells per embryo are indicated on the top right of images. Scale bar: 20µm; n: number of embryos; FC: fold change.

Since PI3K/AKT pathway plays major roles in cell survival and proliferation, the diminution of NANOG-positive cells in LY-treated embryos could result from selective elimination or reduced proliferation of NANOG-positive inner cells. To address this possibility, we first compared the number of mitotic and fragmented nuclei in fixed LY-treated compared to control (vehicle-treated) embryos. As shown in Figure S2F, no differences were observed between the two types of embryos regardless of the time window used for LY treatment. We then performed live imaging of E2.75 H2B-GFP-expressing embryos treated or not with LY for 24 hours. H2B-GFP allowed tracking of mitosis and assessing nuclear fragmentation as a mark of cell death over the course of the movies (Supplemental Movie 1). Control and LY-treated embryos were indistinguishable in terms of proliferation and cell death (Figure S2G-H). We thus conclude that while LY treatment impairs cell survival after Epi and PrE specification, it has no impact on cell survival or cell division at morulae and early/mid-blastocyst stages. Hence, the effects observed for NANOG might be due to direct or indirect regulation of its expression.

Finally, we investigated whether ICM progenitors that express no or low NANOG levels following PI3K inhibition were irreversibly restricted to PrE fate. To that end, we first incubated E2.75 embryos with LY for 24 hours, fixed some embryos to verify the efficacy of the treatment and released the remaining embryos from the inhibitor for an additional 16 hours (Figure 2E). We found no differences between embryos initially treated with the inhibitor or the vehicle, both showing NANOG-only Epi progenitors segregating from GATA4-only PrE progenitors (Figure 2F). Hence, PI3K is strictly required to reversibly maintain NANOG levels rather than to induce its expression. We were unable to assessed whether prolonged incubation with the inhibitor would lead to irreversible extinction of *Nanog* expression since, as we previously reported, LY treatment of late blastocysts compromised inner cell survival (Bessonnard et al., 2019).

We conclude that PI3K maintains NANOG levels by a proliferation and survival-independent mechanism specifically during the early phase of lineage specification when Epi progenitors are first established.

### PI3K inhibition affects Epi and PrE specification

Having established that LY treatment strongly impacts NANOG levels during blastocyst formation, we analysed the effects of PI3K inhibition on the expression of additional lineage-specific TFs. SRY-Box Transcription Factor 2 (SOX2) and Estrogen related receptor beta (ESRRb) are present in all ICM progenitors and become progressively restricted to Epi lineage (Okamura et al., 2019; Wicklow et al., 2014). Similar to NANOG, SOX2 levels in inner cells were significantly reduced after 24 hours of PI3K inhibition (Figure 3A-B). However, SOX2 was less affected after shorter duration of inhibition (Figure S3A). Similar results were obtained for ESSRB (Figure S3B). In contrast, expression levels of POU Class 5 Homeobox 1 (POU5F1/OCT4) and Kruppel Like Factor 4 (KLF4) pluripotency TFs, that become restricted to Epi subsequently (Guo et al., 2010; Morgani and Brickman, 2015), were significantly increased upon PI3K inhibition (Figure S3D-F). The expression of other pluripotency TFs than NANOG likely explains the reversibility of LY treatment and the capacity of cells having temporarily lost NANOG to resume Epi specification upon LY withdrawal.

**Figure 3:**
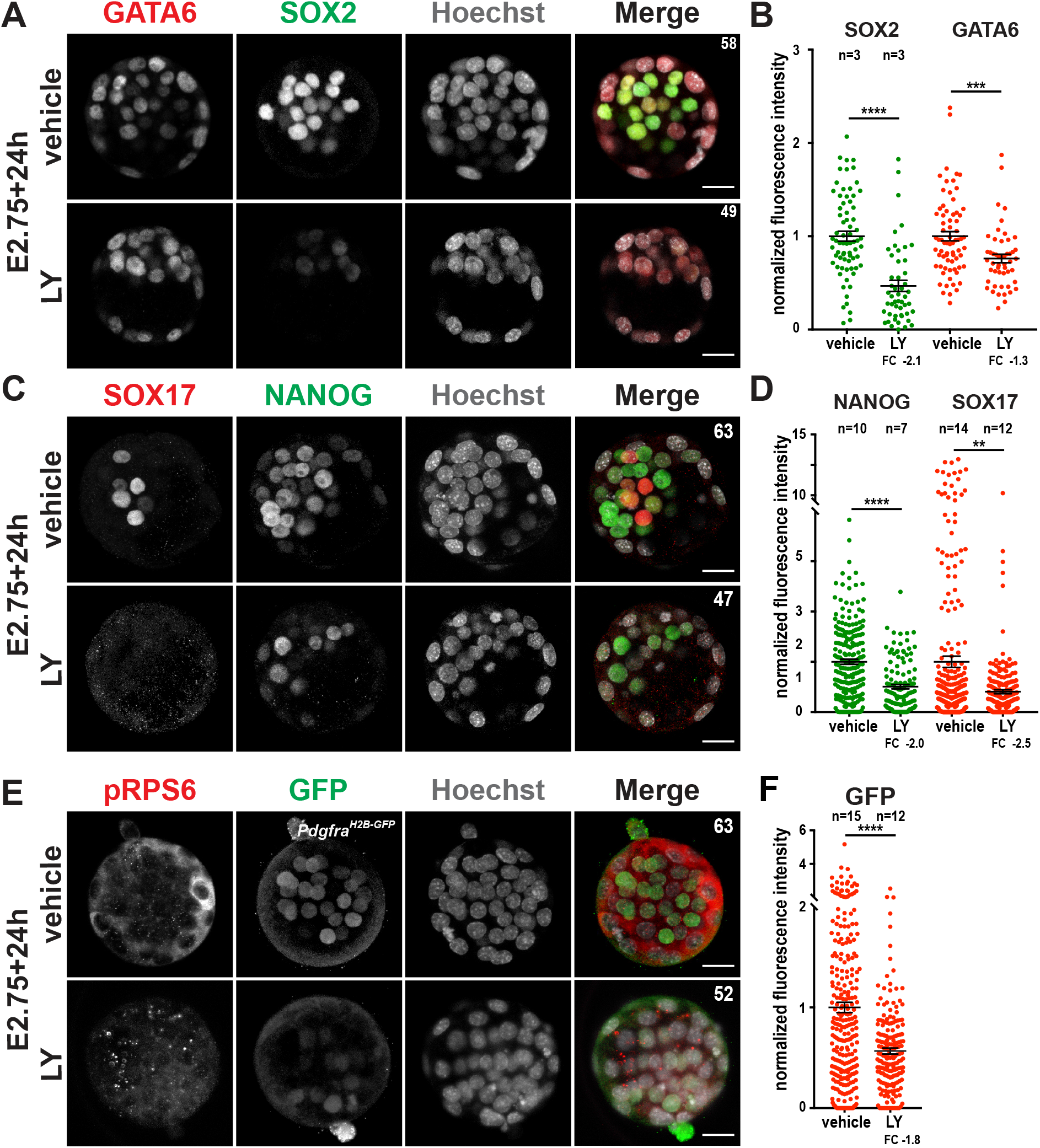
PI3K inhibition affects both Epi and PrE TFs expression. (A, C, E) Immunodetection and (B, D, F) quantification of GATA6 (red), SOX17 (red), pRPS6 (red), SOX2 (green), NANOG (green) and GFP (green) in embryos cultured for 24h from E2.75. Data are from one experiment, representative of 4 (A-B) or combined from two independent experiments (C-F). Pictures do correspond to the projection 5 optical slices. Number of cells per embryo indicated on the top right of images. Scale bar: 20µm; n: number of embryos; FC: fold change.

In absence of NANOG or SOX2, PrE differentiation is compromised as a result of decreased *Fgf4* expression (Frankenberg et al., 2011; Wicklow et al., 2014). Hence, *Nanog-* and *Sox2-*deficient E3.75 blastocysts fail to upregulate platelet-derived growth factor receptor alpha (PDGFRA) and to activate SRY-Box transcription factor 17 (SOX17) and GATA4 binding protein 4 (GATA4) which are normally expressed in a sequential and ERK-dependent manner in PrE progenitors (Artus et al., 2011). Since both NANOG and SOX2 levels are significantly reduced upon PI3K inhibition, we expected a delayed or defective PrE differentiation in LY treated embryos. Accordingly, we observed that while SOX17 expression was detected in multiple ICM cells in vehicle treated embryos, SOX17-positive ICM cells were rarely present in E2.75 embryos treated with LY for 24h (Figure 3C-D). We next looked at *Pdgfrα^H2B-GFP^* expression which is initially low in all ICM cells and increased in PrE progenitors (Plusa et al., 2008). Similar to SOX17, *Pdgfrα^H2B-GFP^* expression was reduced in presence of LY (Figure 3E-F), suggesting that PI3K inhibition prevented ICM differentiation into PrE. Of note, the expression of the key transcriptional TE drivers, Caudal type homeobox 2 (CDX2) and GATA binding protein 3 (GATA3), were hardly affected by LY treatment (Figure S3G-H).

Collectively, our data show that PI3K controls the protein levels of key TFs of the pluripotency network as well as early events of PrE differentiation.

### PI3K inhibition prevents inner cells transcriptome maturation

We next investigated the consequences of PI3K inhibition on gene expression. First, ICMs were immunosurgically isolated from E2.75 embryos treated for 16 hours (∼30-38 cells) and gene expression was analyzed by RT-qPCR on individual ICMs. While both NANOG and SOX2 proteins were downregulated at this time point (Figure 2A-B, 3A-B), only *Nanog* mRNA levels were reduced upon LY treatment (Figure 4A) suggesting different mode of regulation of these two pluripotency TF by PI3K signalling. Notably, *Fgf4* mRNA levels were also reduced in LY-treated ICMs, a likely consequence of the reduced levels of NANOG and SOX2 as suggested by previous studies (Frankenberg et al., 2011; Wicklow et al., 2014). Interestingly, ICM progenitor markers (*Sox21, Wnt7b)* (Allègre et al., 2022; Boroviak et al., 2015; Goolam et al., 2016) were expressed at slightly higher levels in LY-treated ICMs (Figure 4A). Conversely, PrE progenitor markers (*Gata6, Sox17, Pdgfra*) showed a tendency for downregulation (Figure 4A). Together, these data suggest that ICM specification is impaired following PI3K inhibition.

**Figure 4:**
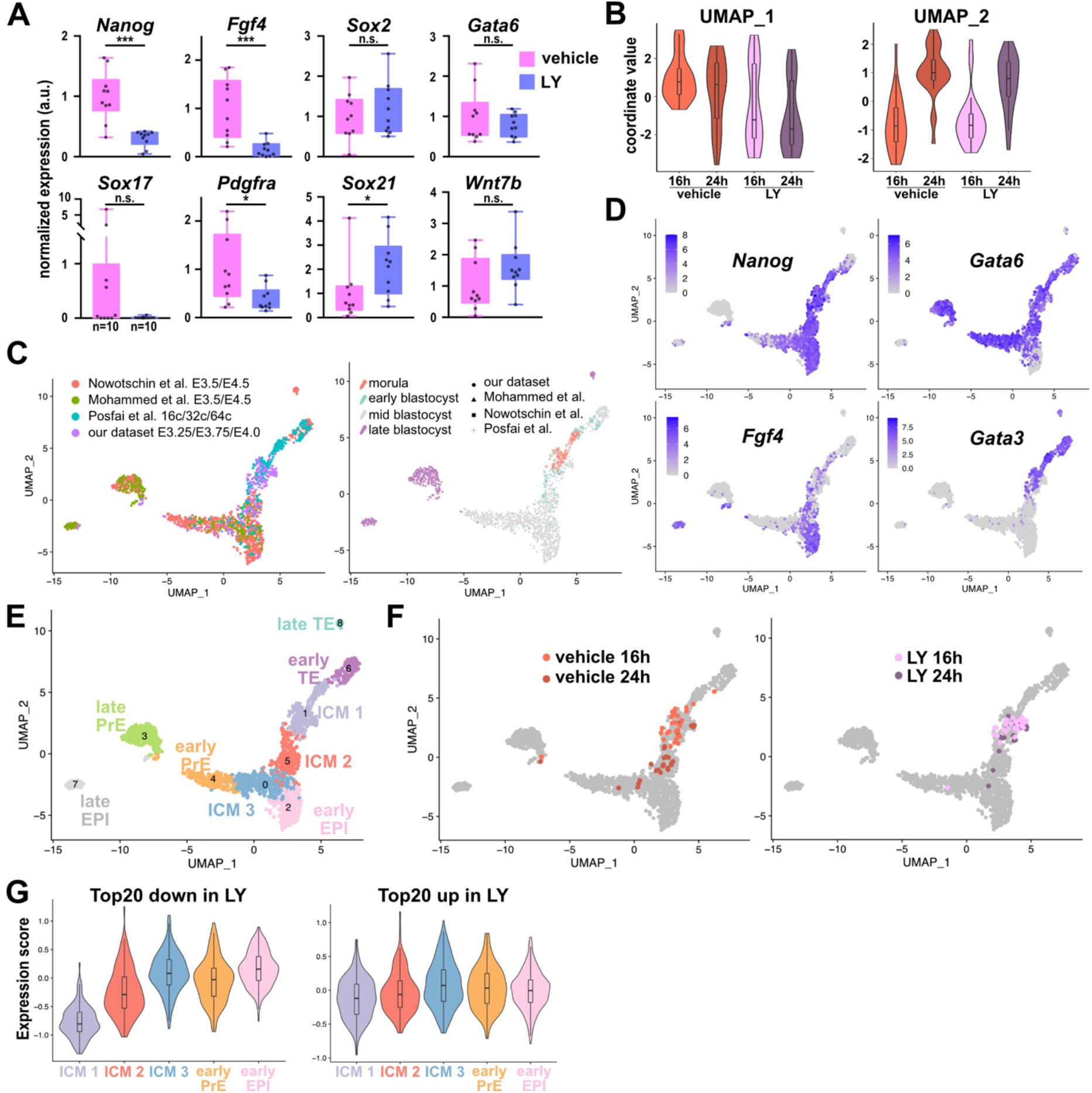
PI3K inhibition prevents the transcriptional changes associated with ICM maturation. (A) Expression of ICM, Epi and PrE markers measured by RT-qPCR on isolated ICM from vehicle-treated (pink) and LY-treated (blue) E2.75 embryos for 24h. (B) Violin plots of coordinated values of vehicle and LY-treated cells along first (UMAP_1) and second (UMAP_2) dimension of the UMAP. (C) UMAP of all integrated cells showing lineage progression. Color indicates the source of the dataset (left) or embryonic stage (right). Symbols indicate the source of the dataset (right). (D) UMAPs of all cells colored by gene expression level of lineage-specific marker. Grey to purple scale indicates average expression. (E) UMAP showing the 9 identified clusters. (F) UMAP of all integrated single cells. Single cells from external datasets in grey (unselected), vehicle (left) and LY-treated (right) single inner cells from our dataset colored by duration of culture. (G) Violin plots of the average expression score in clusters 1, 5, 0, 2 and 4 of the Top20 most downregulated (left) and upregulated (right) genes after LY treatment.

To obtain a comprehensive view of the impact of LY treatment on the transcriptome of inner cells, single-cell RNA sequencing (scRNAseq) was performed on manually dissociated ICM recovered from E2.75 embryos, after treatment for 16 and 24 hours with vehicle or LY (31× E2.75+16h vehicle, 31× E2.75+16h LY, 28× E2.75+24h vehicle, 24× E2.75+24h LY). We applied uniform manifold approximation and projection (UMAP) dimensionality reduction and found that cells separated according to treatment and duration (UMAP_1 and _2 respectively, Figure 4B and S4A). To better appreciate the effect of LY treatment on the inner cell transcriptome, we compared our treated cells data to datasets from non-cultured embryos at the morula/blastocyst stages: 20 inner cells from E3.25, 22 inner cells from E3.75 and 14 inner cells from E4.0 blastocysts generated in this study as well as published scRNA-seq datasets from 16-, 32- and 64-cell stage embryos (262 cells) (Posfai et al., 2017), and from E3.5 and E4.5 embryos (1083 cells) (Mohammed et al., 2017; Nowotschin et al., 2019). UMAP dimensionality reduction of the combined datasets showed extended mixing of cells from the different studies, reflecting developmental lineage progression in accordance with lineage-specific markers expression (Figure 4C-D and S4B). Clustering analysis identified 9 clusters which, based on their gene expression profile, distinguished inner cells and ICM progenitors (clusters 1, 5 and 0), Epi (clusters 2 and 7), PrE (clusters 4 and 3) and outer cells and TE (clusters 6 and 8) differentiation (Figure 4E, S4B-C and Supplementary Table 1). Interestingly, vehicle- and LY-treated cells were distributed in a somewhat different patterns on the UMAP: most LY-treated cells mapped to the ICM progenitors cluster 1, regardless of culture duration, while more than two thirds of 24h vehicle-treated cells were found in more mature clusters (6, 0, 3 and 4) (Figure 4F). Accordingly, the expression of the top 20 genes downregulated in LY-treated cells increased during ICM specification (Figure 4G and supplementary table 2). Altogether, our analyses suggest that LY treatment blocks the normal developmental progression of the ICM.

### GSK3β is a downstream effector of PI3K/AKT regulating NANOG levels in ICM progenitors

Next, we searched for the effectors acting downstream of PI3K and potentially involved in the regulation of Epi and PrE TFs. First, we addressed the role of mTOR using mTORC1 (Rapamycin) and dual (INK128) inhibitors. Inhibition of mTORC1 and mTORC2 signalling did not alter NANOG expression (Figure 5A-B and S5A-B) indicating that mTOR is not acting downstream of PI3K/AKT to maintain NANOG levels. FOXO TFs are downstream direct targets of PI3K/AKT, which translocate to the cytosol upon AKT-mediated phosphorylation (Brunet et al., 1999). Importantly, FOXO1 and FOXO3 proteins have been shown to regulate pluripotency in human and mouse ES respectively, possibly through direct transcriptional regulation of *Sox2 and Pou5f1/Oct4* pluripotency genes (Zhang et al., 2011). Both nuclear and cytoplasmic FOXO3 staining were observed in control blastocysts with some cells showing nuclear accumulation and others nuclear exclusion (Figure S5C). Such heterogeneous nucleocytoplasmic distribution may reflect the heterogenous PI3K/AKT activity observed at this stage (Figure 1). Accordingly, we observed a clear FOXO3 nuclear accumulation in most cells of LY-treated embryos (Figure S5C). Assuming that the positive transcriptional regulation of FOXO proteins on pluripotent Epi TFs is not only operational in ES cells but also *in vivo* in the embryo, this suggests that FOXO3 is not the main direct effector acting downstream of PI3K/AKT in ICM progenitors to positively regulate SOX2 and NANOG levels. Another well-known target of AKT is the glycogen synthase kinase 3 (GSK3) α/β, the activity of which is inhibited upon AKT phosphorylation on Ser31 and Ser9 respectively (Cross et al., 1995). In ES cells, inhibition of GSK3 α/β activity by PI3K/AKT has been proposed to participate to NANOG upregulation and maintenance of pluripotency (Niwa et al., 2009; Paling et al., 2004; Singh et al., 2012). Accordingly, E2.75 morulae treated for 24h with the GSK3 inhibitor CHIR99021 (CHIR) showed a significant upregulation of NANOG levels (Figure 5C-D). This indicates that NANOG levels in ICM progenitors are regulated by GSK3 α/β activity. *Gsk3β* is more highly expressed than *Gsk3α* in ICM cells (Boroviak et al., 2015; Deng et al., 2014). We thus analyzed the phosphorylation status of GSK3β on Ser9 in E2.75 embryos cultured for 24h. Similar to pRPS6, heterogeneous pSer9-GSK3β levels were observed in both TE and ICM cells (Figure 5E-F). Importantly, LY treatment downregulated pSer9-GSK3β levels, although mitotic cells generally retained some pSer9-GSK3β signal (Figure 5E and S5D). Collectively, our data suggest that PI3K/AKT is acting through GSK3β kinase to regulate NANOG levels in ICM progenitors.

**Figure 5:**
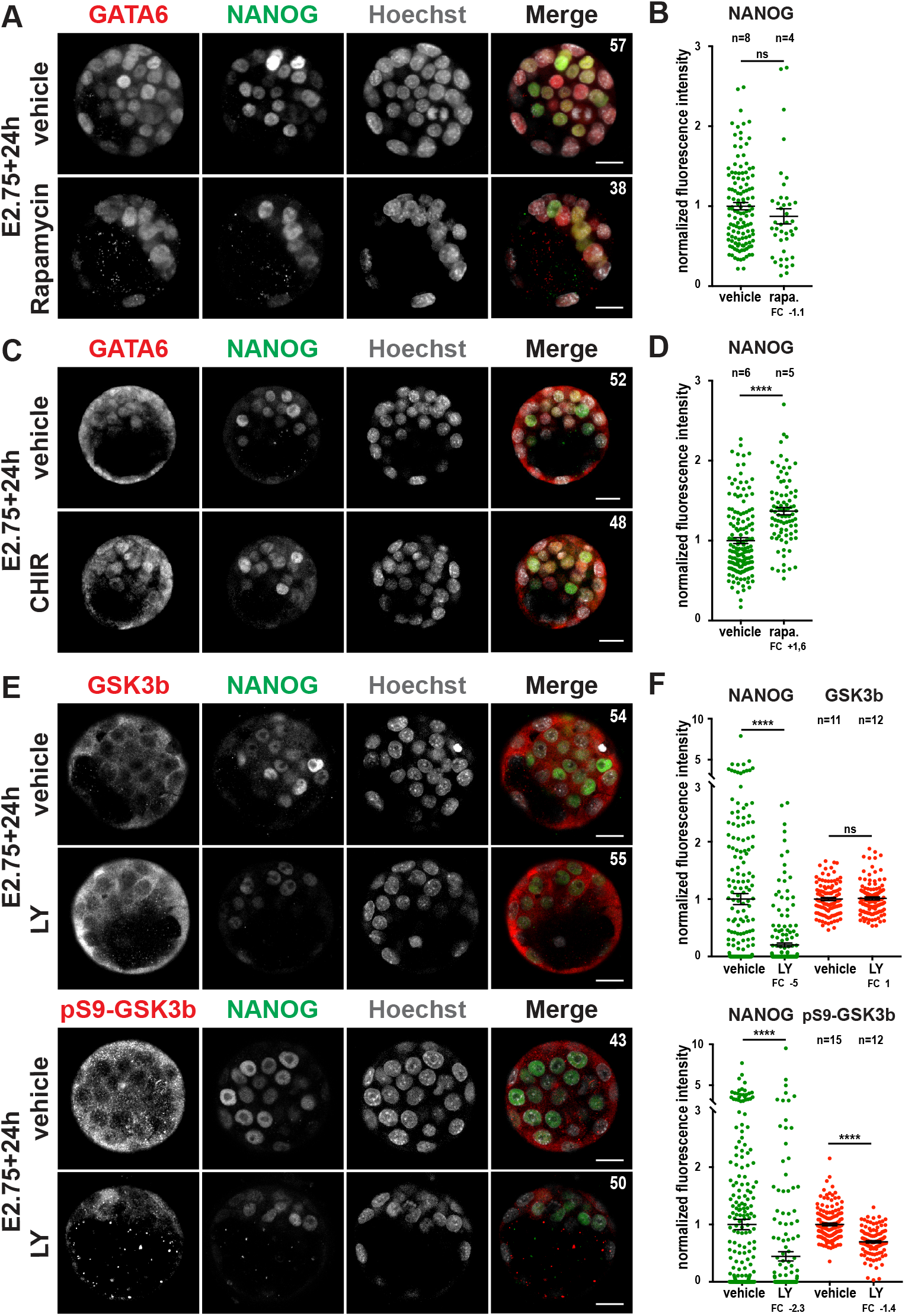
GSK3 is a downstream target of PI3K/AKT that regulates NANOG levels in ICM progenitors. (A) Immunodetection of GATA6 (red) and NANOG (green) and (B) quantification of NANOG (green) in embryos cultured with rapamycin for 24h from E2.75. (C) Immunodetection of GATA6 (red) and NANOG (green) and (D) quantification of NANOG (green) in embryos cultured for 24h with CHIR99021 from E2.75. (E) Immunodetection of panGSK3b (red), phosphoSer9-GSK3b (red) and NANOG (green) and (F) quantification of panGSK3b (red), phosphoSer9-GSK3b (red) and NANOG (green) in embryos cultured with LY for 24h from E2.75. Data are from one experiment, representative of two to three independent experiments. Pictures do correspond to the projection of 5 optical slices. Number of cells per embryo indicated on the top right of images. Scale bar: 20µm, n: number of embryos; FC: fold change.

### PI3K regulates NANOG independently of ERK and confers ICM progenitors responsiveness to FGF4

As discussed, the main drivers of *Nanog* downregulation during ICM to PrE progenitor differentiation are GATA6 and ERK (Bessonnard et al., 2014; Hamazaki et al., 2006; Santostefano et al., 2012; Schrode et al., 2014; Tosenberger et al., 2017). We thus addressed whether downregulation of NANOG upon PI3K inhibition required either of these factors. In *Gata6* mutant blastocysts, NANOG is expressed in all ICM cells (Bessonnard et al., 2014; Schrode et al., 2014). Interestingly and contrary to their wild-type and heterozygous littermates, zygotic (Figure S6C) as well as maternal and zygotic (Figure 6A) *Gata6* homozygous mutants displayed robust NANOG nuclear staining in most inner cells following LY treatment. Quantification confirmed that NANOG levels were significantly higher in *Gata6* mutant embryos (Figure S6A-B). GATA6 seems therefore required for efficient downregulation of NANOG upon PI3K inhibition. Whether PI3K acts through GATA6 to modulate NANOG levels or whether this is due to higher NANOG levels in ICM progenitors of *Gata6-/-* embryos irrespective of PI3K activity (Bessonnard et al., 2014) remains to be determined.

**Figure 6:**
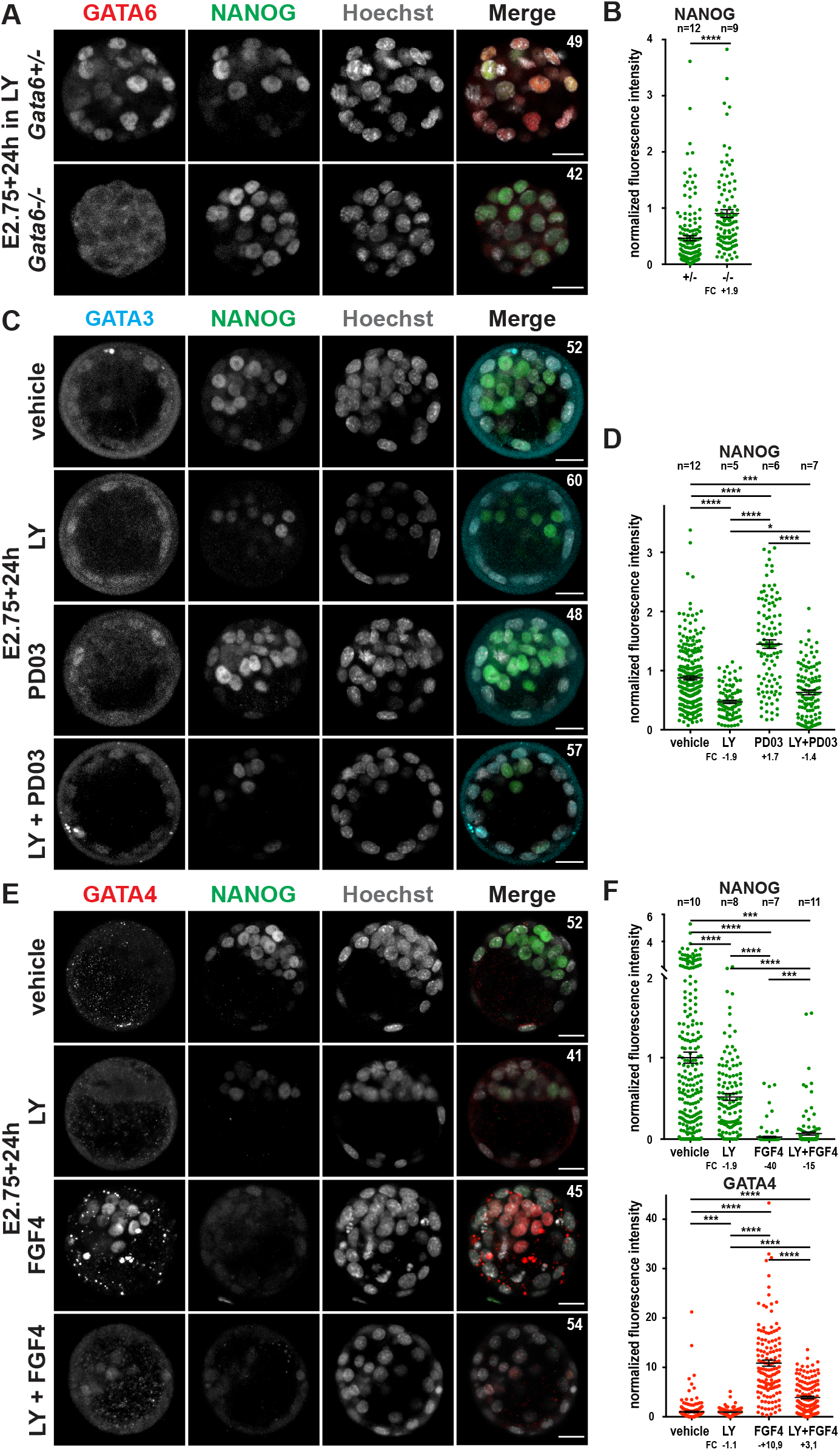
PI3K regulates NANOG levels independently from ERK and confers ICM progenitors responsiveness to FGF4. (A) Immunodetection of GATA6 (red) and NANOG (green) and (B) quantification of NANOG in maternal-zygotic *GATA6 +/- and -/-* embryos cultured for 24h from E2.75. (C, E) Immunodetection of GATA3 (cyan), GATA4 (red) and NANOG (green) and (D, F) and (B) quantification of NANOG and GATA4 in embryos cultured for 24h from E2.75 with various compounds. Graphs shows results from two (F) or three (B, D) independent experiments. Pictures do correspond to the projection of 5 optical slices. Number of cells per embryo indicated on the top right of images. Scale bar: 20µm; n: number of embryos; FC: fold change.

Then, we addressed the possibility that PI3K/AKT may regulate ICM differentiation through direct interactions with the FGF/ERK pathway. Indeed, similar to PI3K inhibition, the addition of high concentration of FGF4 and heparin during the period of Epi/PrE specification is sufficient to repress NANOG and SOX2 (Wicklow et al., 2014; Yamanaka et al., 2010). Together with the previously identified crosstalks between PI3K/AKT and RAF/MEK/ERK pathways (Mendoza et al., 2011; Zimmermann and Moelling, 1999), this raised the possibility that the effect of LY on NANOG is mediated, at least in part, by MEK/ERK. We asked whether the impact of PI3K inhibition was modified by the simultaneous inhibition of MEK. Consistent with previous reports (Bessonnard et al., 2017; Saiz et al., 2016; Yamanaka et al., 2010), E2.75 embryos treated for 24h with the MEK inhibitor PD03 exhibited robust NANOG expression in all ICM cells (Figure 6C-D). In contrast, the simultaneous inhibition of PI3K and MEK led to NANOG downregulation. Thus, PI3K inhibition affects NANOG levels in ICM progenitors independently of MEK/ERK activity, and strikingly the effect of PI3K is dominant.

Finally, we asked whether defective PrE differentiation of LY-treated embryos could be rescued by exogenous FGF4. We compared E2.75 embryos cultured for 24h with FGF4 and heparin alone or in combination with LY. As expected, exposure to exogenous FGF4 promoted NANOG downregulation and GATA4 activation in most ICM cells (Figure 6 E-F). Strikingly, when PI3K was inhibited, GATA4 activation by exogenous FGF4 was largely blocked (Figure 6 E-F). Hence, despite the presence of the two mains drivers of PrE fate, namely GATA6 and FGF4, ICM progenitors treated with LY are unable to engage into PrE differentiation. Thus, our data demonstrate that PI3K activity is required for ICM progenitor responsiveness to FGF signalling.

## DISCUSSION

In this study, we investigated whether the regulation of NANOG by PI3K that has been described in mouse ES cells was also operating during the establishment of the pluripotent Epi in the mouse embryo. We showed that PI3K/AKT is active during preimplantation development and positively controls NANOG levels in ICM progenitors during the early phase of Epi/PrE specification. PI3K signalling sustains the expression of other Epi TFs such as SOX2 and ESSRb but not that of OCT4 and KLF4. We further showed that PI3K is not acting through MEK/ERK to modulate the levels of these TFs. Our results therefore establish PI3K signaling as a new important player in the regulation of key Epi TFs in the mouse blastocyst. At the early stages of ICM differentiation, when FGF/ERK signaling is less prominent, positive regulation of pluripotency TFs by sustained PI3K activity in ICM progenitors may bias the activity of the gene regulatory network towards a robust pluripotency network driving the definitive acquisition of the Epi fate. When Epi-derived FGF4 concentration increases within the ICM, PI3K activity is then required in the remaining ICM progenitors to respond efficiently to FGF signalling and engage into PrE fate (Figure 7).

**Figure 7:**
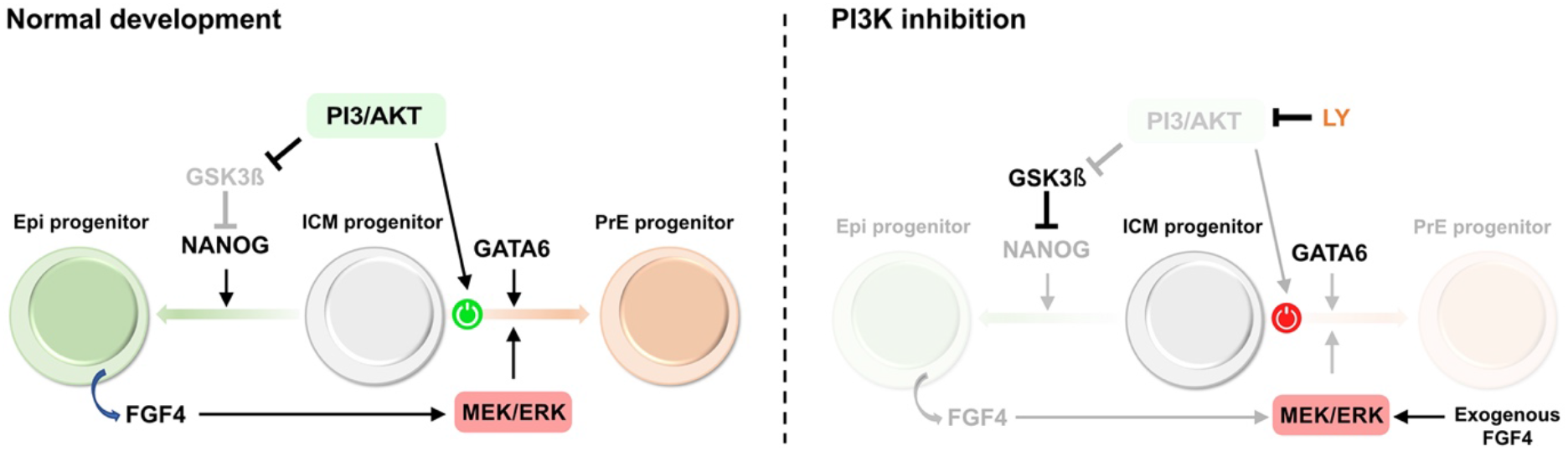
Proposed model of ICM progenitor specification. The ICM progenitor state is characterized by the co-expression of the antagonistic TFs NANOG and GATA6. The activity of GSK3β, a negative regulator of NANOG, is regulated by PI3K/AKT activity. NANOG, together with other factors, promotes the reinforcement of the pluripotency TFs network leading to Epi specification. Epi progenitors secrete FGF4 that, with GATA6, will drive the MEK/ERK-driven differentiation of surrounding ICM progenitors into PrE. PI3K/AKT activity is also required for ICM progenitor to engage into PrE differentiation. In absence of PI3K activity (right), GSK3β activity is upregulated leading to NANOG downregulation. Consequently, Epi differentiation is blocked, local concentration of FGF4 in the ICM remains low and PrE differentiation is impaired. In presence of exogenously provided FGF4, ICM progenitor is unable to respond to the inductive signal and PrE differentiation is not rescued.

Epi/PrE specification is a rapid process that occurs in a few hours and involves the downregulation of either NANOG or GATA6 followed by the reinforcement of the pluripotency program in Epi progenitors, versus the sequential activation of lineage-specific TFs in PrE progenitors. The initial downregulation of NANOG and GATA6 could result from multiple mechanisms acting both at the transcriptional and post-transcriptional levels. We found that both *Nanog* mRNA and protein levels were affected by PI3K inhibition. NANOG has a short half-live (1-3 hours compared to 6h or more for OCT4 in ES cells (Sanchez-Ripoll et al., 2013)), therefore variations in mRNA expression or translation efficiency rapidly impact NANOG levels. Importantly, PI3K acts upstream of mTORC signalling, and inhibition of this pathway in mouse blastocysts or ES cells, induces a paused state characterized by a global downregulation of transcription and translation (Bulut-Karslioglu et al., 2016; Xu et al., 2021). In this study, we showed that inhibition of mTORC signaling during the early phase of ICM specification did not alter NANOG levels indicating that NANOG downregulation following PI3K inhibition was not simply a consequence of entering into mTORC-mediated “paused” state. Contrary to *Nanog, Sox2* mRNA levels were hardly affected by LY treatment while SOX2 protein levels were significantly reduced. PI3K can modulate protein levels through AKT-mediated phosphorylation as AKT has been shown to directly phosphorylate SOX2, OCT4 and KLF4 and modulate their stability (Chen et al., 2013; Fang et al., 2014; Lin et al., 2012). Whether SOX2 half-life is regulated by PI3K/AKT in ICM progenitor will require further investigations.

PI3K has been shown to regulate pluripotency in ES cells by additional mechanisms including through FOXO TFs. However, both the subcellular localization of FOXO3 and the lack of a major transcriptional response in LY-treated embryos argue against such mechanism having a prominent role in the embryo in ICM progenitors. In contrast, our data suggest that GSK3 acts downstream of PI3K/AKT to regulate NANOG. In ES cells, GSK3 activity has been shown to negatively regulate *Nanog* through either stabilization of TCF7L1, which together with β-catenin represses the transcription of *Nanog* (Wray et al., 2011; Yi et al., 2011) or through destabilization of c-MYC, which promotes pluripotency and represses PrE program (Bechard and Dalton, 2009; Cartwright et al., 2005; Singh et al., 2012; Smith et al., 2010). Through AKT-mediated inhibitory phosphorylation, continuous PI3K signalling could help to keep in check GSK3 activity and thereby to maintain NANOG levels in ICM progenitors. GSK inhibition has also been shown to enhance *Nanog* mRNA translation in ES cells (Sanchez-Ripoll et al., 2013). In the future, it will be important to determine the respective contribution of these mechanisms in the regulation of NANOG levels in ICM progenitors.

Both *Gata6* and FGF/ERK signalling negatively regulate *Nanog* expression in embryo and ES cells. We found that NANOG downregulation following PI3K inhibition was less efficient in *Gata6* mutant embryos suggesting that PI3K regulates NANOG at least in part through GATA6. In contrast, we showed that PI3K is not acting through ERK: contrary to other cells types including ES cells (Mendoza et al., 2011; Paling et al., 2004), PI3K/AKT and RAS/ERK pathways do not seem to be interconnected in ICM progenitors, at least in regulating NANOG. Independence from ERK signalling was further confirmed by the fact that NANOG downregulation occurs in absence of ICM progenitor maturation and *Fgf4* upregulation. Interestingly, no impact on NANOG levels was observed when PI3K was inhibited at later stage (E3.25, ∼50 cells). At that time, coordinated expression of pluripotency markers has been established (Allègre et al., 2022) which may render ICM cells more resistant to perturbation in the levels of single pluripotency TFs. When applied at E2.75, PI3K inhibition may instead delay the establishment of such coordinated expression and consequently postpone ICM maturation. An alternative but non-exclusive explanation is that the control of TFs levels by PI3K is temporally regulated.

Activation of the PrE-specific genetic programme in ICM progenitors is triggered by FGF/ERK signalling and requires GATA6 (Bessonnard et al., 2014; Chazaud et al., 2006; Kang et al., 2013; Kang et al., 2017; Molotkov et al., 2017; Nichols et al., 2009; Schrode et al., 2014; Yamanaka et al., 2010). Here, we show that this is not sufficient as LY-treated ICM progenitors expressing GATA6 at normal levels do not properly activate the PrE programme when exposed to FGF4. Therefore, PI3K activity is required to confer ICM progenitors with the competence to respond efficiently to PrE-inductive cues. In the embryo, Epi progenitors are rapidly committed once specified and, despite sustained levels of FGFR1, they do not adopt a PrE fate in response to FGF4 (Bessonnard et al., 2017; Kang et al., 2017; Molotkov et al., 2017; Saiz et al., 2016). Thus, within the same microenvironment, specified Epi and uncommitted ICM progenitors respond differently to FGF4. Here, we found that inner cells display disparate PI3K activity by the time of ICM specification. It is therefore possible that transient downregulation of PI3K activity may help, alongside other mechanisms (Azami et al., 2019; Kang et al., 2017), to prevent Epi progenitor from changing fate under the action of autocrine/paracrine FGF4 activity. That PI3K activity may act as a permissive signal during ICM specification is an attractive hypothesis that will require to dynamically monitor PI3K activity in the embryo and determine how its modulations in ICM progenitors correlate with cell fate acquisition.

In conclusion, our study unravels a dual role of PI3K/AKT signaling: this pathway both maintains expression of the pluripotent TFs NANOG, ESRRb and SOX2 and safeguards the competence to response to FGF signaling at the onset of Epi and PrE specification. Although PI3K/AKT and GSK3 have been long known to be critical determinants of pluripotency in mouse ES cells, evidence for PI3K/AKT regulating early embryonic fate was lacking so far. There are many reasons for this, in particular the fact that the role of PI3K seems to be critical only before the establishment of a robust pluripotency network and the rise of the FGF/ERK signalling pathway. The facts that specification events occur very rapidly in the mouse preimplantation embryo and that the PI3K/AKT pathway regulates proliferation and survival once lineages are established has also delayed the discovery of the initial role of PI3K in regulating ICM progenitor fate. Our findings will be instrumental to re-evaluate the role of PI3K/AKT signalling during cell fate decisions in other mammalian embryos. Indeed, recent observations in pig (Ramos-Ibeas et al., 2019) and human (Wamaitha et al., 2020) preimplantation embryos suggest that regulation of *Nanog* by PI3K may be conserved across species.

## AKNOWLEDGMENTS

We are grateful to the staff of the animal facility of Institut Pasteur for animal care and their help during this work. We thank Jean-Yves Tinevez from the Image Analysis Hub of Institut Pasteur for his help with the Icy software. We thank Jérôme Artus, Claire Chazaud and Nicola Festuccia for critical reading of the manuscript, Maud Borensztein and Raquel Pérez-Palacios for their advice on single cell cDNA and library preparation, Nicola Festuccia for his help in single ICM RT-qPCR and Luis Fernando Altamirano Pacheco for scRNAseq data processing. This work was supported by the Institut Pasteur, the Centre National de la Recherche Scientifique and the Agence Nationale de la Recherche (ANR-10-LABX-73-01 REVIVE). A.G. was supported by Sorbonne Université and received fellowship from the French Ministère de l’Enseignement Supérieur et de la recherche, the Fondation ARC pour la Recherche sur le Cancer and the REVIVE Labex. A. M. was supported by Université Paris Cité and received fellowship from the French Ministère de l’Enseignement Supérieur et de la Recherche.

## Competing interests

The authors declare no competing interest.

## AUTHORS CONTRIBUTIONS

A.G., Conception and design, Acquisition of data, Analysis and interpretation of data, Drafting or revising the article; A.M., Acquisition of data, Analysis and interpretation of data, Drafting or revising the article; S.V.-P., Acquisition of data; V.L., S.M. Analysis and interpretation of data; P.N., Analysis and interpretation of data, Drafting or revising the article; M.C.-T., Conception and design, Analysis and interpretation of data, Drafting or revising the article.

## Data availability

scRNA-seq data supporting the findings of this study have been deposited in NCBI GEO.

## MATERIALS AND METHODS

### Embryo collection and culture

All experiments were performed according to the French and European regulations on care and protection of laboratory animals (EC Directive 86/609, French Law 2001-486 issued on June 6, 2001) and were approved by the Institut Pasteur ethics committee. All mice were kept in a 14h light cycle from 7am to 9pm, except mice used to obtain E3.0 embryos which were kept in a 14h light cycle from 2pm to 1am. Preimplantation embryos were obtained from natural mating of CD1 mice (Charles River Laboratories, France). Embryos were staged according to the date of the vaginal plug (E0.5) at noon and collected by flushing oviducts or uteri with FHM medium (Millipore). Embryos were cultured at low density in KSOM+AA medium (Millipore) using 4-well plates (Nunc) at 37°C, 8% CO2. The solvent which was used to resuspend the molecules was added to the culture containing the control embryos. Compounds used in this study are listed in Supplementary Table 3.

### Mutant mice and genotyping

Zygotic *Gata6^+/−^* mice were the result of mating *Gata6^tm2.2Sad^* males (Sodhi et al., 2006) with Tg(Pgk1-cre)1Lni females (Lallemand et al., 1998). By crossing *Gata6^+/−^* mice, 25% of the obtained embryos were *Gata6^-/−^*. To achieve maternal-zygotic deletion of *Gata6*, we first crossed female *Zp3Cre^Tg/Tg^* mice (Vries et al., 2000) with *Gata6^flox/flox^* males (Sodhi et al., 2006) to produce *Zp3Cre^Tg/0^;GATA6^flox/+^* females. Crossing these females with CD1 males, led to Cre-recombination of *Gata6* in 100% of the offspring and we used the resulting *Zp3Cre^Tg/0^;GATA6^+/del^* males for the final mating. We also crossed *Zp3Cre^Tg/Tg^* female with *Zp3Cre^Tg/0^;GATA6^flox/+^* males to produce *Zp3Cre^Tg/Tg^;GATA6^flox/+^* males and females that were then intercrossed to obtain *Zp3Cre^Tg/Tg^;GATA6^flox/del^* females. Mating of *Zp3Cre^Tg/Tg^;GATA6^flox/del^* females with *Zp3Cre^Tg/0^;GATA6^+/del^* males resulted in the maternal-zygotic deletion of *Gata6* in 50% of the embryos. For live imaging experiments, heterozygous *CAG::H2B-EGFP* males (Hadjantonakis and Papaioannou, 2004) were crossed with female CD1 to obtain H2B-EGFP expressing embryos with a frequency of 50%. Embryo genotyping was performed after confocal imaging by incubating single embryos in lysis buffer (10mM TrisHCl ph8; 50mM KCl; 0.01% gelatine; 300µg/ml Proteinase K (Thermo Fisher)) at 56°C for 1h. Proteinase K was inactivated at 95°C for 10 minutes. The amplification program for both steps was as follows: 30 sec 98°C followed by 25 cycles of 10 sec 98°C, 15 sec 61°C and 1 min 72°C, and then 3 min 72°C. Primers used were G6-del-F (AGTCTCCCTGTCATTCTTCCTGCTC), G6-del-R (TGATCAAACCTGGGTCTACACTCCTA) and G6-WT-F (GTGGTTGTAAGGCGGTTTGT). Amplicon sizes are 568 bp, 250 b and 159 bp for *GATA6^flox^, GATA6^del^* and *GATA6^+^* alleles respectively.

### Whole-mount immunostaining

For immunofluorescence stainings, embryos were fixed either with 4% paraformaldehyde (PFA) in PBS for 15 minutes at room temperature (RT) or, for pRPS6 and pS9-GSK3b stainings, with 8% PFA in PBS supplemented with phosSTOP (Roche) for 1 hour at 4°C. Embryos were then washed three times with 0.1% PBS Tween, permeabilized with PBS 0.1% Triton-X100 (PBX) for 30 minutes at RT and incubated in blocking solution (10% donkey serum in PBX) for 1h. Embryos were treated with primary antibodies diluted in blocking solution at 4°C overnight. After three washes with PBX, embryos were incubated with secondary antibodies diluted 1/300 in PBX for at least 1h. Nuclear staining was achieved by Hoechst 33342 (Thermo Fisher). Primary and secondary antibodies used in this study are listed in Supplementary Table 4.

### Confocal microscopy

All images were acquired by using Zeiss LSM800 and LSM900 confocal microscopes. Embryos were placed in an in-house developed and produced eggbox imaging device (Vandormael-Pournin et al., 2021) to facilitate a multi-positioning setup and scanned using a plan-apochromat 20× objective without immersion. Images were acquired using bi-directional scanning, 2-line averaging, 2× zoom and 1 airy unit pinhole. Stacks were created with a 2µm step, generating 16-bit (512×512px) images. For live imaging, embryos were imaged in a 10 min time interval for a total of 24h and maintained in a humified incubation chamber at 37°C in 5% CO_2_.

### Image analysis

ImageJ (NIH) was used to manually perform cell counting and obtain projections of optical slices. Quantification of nuclear fluorescence intensities was achieved using the Spots creation wizard of the IMARIS (Bitplane) software. To correct the z-associated attenuation of fluorescence, nuclear signals were normalized by dividing their values by the corresponding value of Hoechst. Calculation of the background level was performed by dividing the average of the mean fluorescence intensities of randomly chosen cytoplasmic spots with the average of Hoechst fluorescence. Nuclei with mean fluorescence intensities below background level were set to zero. The Icy software (Institut Pasteur) was used to measure mean fluorescence intensities of cytoplasmic proteins. For each cell a 2D region of interest (ROI) was manually drawn in the nucleus and cytoplasm. In order to compare protein levels between different developmental stages, the values of Hoechst were corrected according to the nuclear volume of the corresponding stage before they were used to normalize the measured nuclear and cytoplasmic signals. The correction factors for each stage were obtained from (Aguirre-Lavin et al., 2012). Quantification shown on figure panels 3E, 6E and S6B were obtained after pooling experiments with limited variation in intensity levels across them. Elsewhere, quantifications are from a single experiment representative of at least two independent experiments.

### Statistical analysis

Graphpad Prism software was used to perform statistical tests. Significant correlation was determined by fitting data to linear regression model. The non-parametric Wilcoxon-Mann-Whitney test was applied to assess statistical significance of cell counts and the Kolmogorov-Smirnov test was used to calculate significance of normalized fluorescence intensities. Error bars indicate ± standard error to the mean deviation. Significance was defined as: not significant (ns) ≥ 0.05; *p < 0.05; **p < 0.01; ***p < 0.001; ****p < 0.0001.

### Single embryo RT-qPCR

The following protocol is based on the FLASH-seq protocol (Festuccia et al., 2023; Hahaut et al., 2022). Embryos were cultured and immunosurgery was performed as described below to isolate single ICMs. Single ICMs were individually transferred into PCR tubes containing 5 μL of lysis buffer (0.1 μL Triton X-100 (10% v/v in water, Sigma-Aldrich); 1.2 μL dNTP mix (25 mM each, Thermo Fisher); 0.46 μL 5’Bio-AAGCAGTGGTATCAACGCAGAGTACT30VN-3’ (100 μM, IDT); 0.46 μL 5’ Biotin-AGTGGTATCAACGCAGAGTAC G ATC NNNNNNNN rGrGrG-3′Phospate (100 μM, IDT); 0.15 μL recombinant RNase Inhibitor (40 U/μL, Thermo Fisher); 0.06 μL Dithiothreitol (DTT, 100 mM, Thermo Fisher); 1 μL betaine (5M, Sigma Aldrich); 0.45 μL dCTP (100 mM, Sigma Aldrich); sterile water. Then, PCR tubes were transferred to the thermocycler for 3 min at 72°C and immediately put on ice. The following RT-qPCR mix was added to the tubes: 1.19 μl DTT; 4 μL betaine; 0.23 μL magnesium chloride (1 M, Thermo Fisher); 0.48 μL recombinant RNase Inhibitor; 0.25 L Superscript IV Reverse Transcriptase (200 U/μL, Thermo Fisher); 12.5 μL KAPA HiFi Hot-Start ReadyMix (2×, Roche); sterile water. The following program was run on the thermocycler containing the tubes: 60 min at 50 °C, 90 °C for 3 min, then 20 cycles of (98 °C for 20 s, 67 °C for 20 s, 72 °C for 6 min with the addition of 12s per cycle). PCR products were purified using AMPure XPbeads (Beckman Coulter) and eluted in 15 μl of sterile water. Real-time quantitative PCR performed in duplicates in 384-well plates with a LightCycler 480 (Roche) using SYBR Green I Master (Roche) and 2 µl of 50X dilution of single ICMs sample. - Standard and melting curves were generated to verify the amplification efficiency and the production of single DNA species. Relative concentrations were determined and normalised to the geometric mean of three reference genes (*Hprt, Tcf25* and *Zfp330*) showing minimal variations at morula and early/mid blastula stage according to RNAseq dataset. Primers used in this study are listed in Supplemental Table 5.

### Single cell isolation, library preparation and sequencing

Single inner cells were collected from E2.75 embryos culture for 16h and 24h (∼30-38 cells and ∼∼50-64 cells respectively) as well as non-cultured E3.25 (∼47-55 cells), E3.75 (∼63-69 cells) and E4.0 (∼96-108 cells) blastocysts. Single cells from each stage and condition were obtained from two to three independent experiments, each including multiple litters. For each experiment, some embryos were fixed and analysed by immunofluorescence to verify the development stage of the embryos and the effectivness of LY treatment, while the remaining embryos were processed for low-input RNA-sequencing. The zona pellucida of embryos was removed using acidic Tyrode’s solution (Sigma) and single ICM were isolated by immunosurgery. First, embryos were cultured for 30 minutes at 37°C in anti-mouse red blood cell serum from rabbit (Rockland) and transferred to guinea pig complement serum (Sigma) for 20 minutes at 37°C. The lysed TE cells were carefully removed by mouth pipetting in PBS containing acetylated bovine serum albumin (ac-BSA). To obtain single cells, isolated ICM were briefly exposed to TripLE (Gibco) and then manually dissociated by repeated mouth pipetting with 40µm glass pipettes in PBS/ac-BSA. Single cells were individually picked into 1.5ml Eppendorf tubes, directly transferred to -80°C or immediately processed. Following the protocol of (Tang et al., 2010) and adapted by (Borensztein et al., 2018), mRNAs from every single cell were extracted, reverse-transcribed from 3’UTR and amplified. As a quality check, the amplification yield of the housekeeping genes Gapdh (F-5’-CCCCAACACTGAGCATCTCC-3’, R-5’-ATTATGGGGGTCTGGGATGG-3’) and Hprt (F-5’-CCTGTGGCCATCTGCCTAGT, R-5’-GGGACGCAGCAACTGACATT-3’) was assessed by RT-qPCR using SyBR Green solution on a LightCycler480 (Roche). The cDNAs of every single cell were purified using the DNA Clean & Concentrator Kit (Zymo) and loaded and run on 2% agarose gels. cDNAs larger than approximately 500 bp were excised from the gel and purified using the Zymoclean Gel DNA Recovery Kit (Zymo). The concentration of the purified cDNAs was quantified using the Qubit dsDNA HS Assay Kit (Thermo Fisher). The Nextera XT DNA Library Prep Kit (Illumina) was performed according to manufacturer’s instructions to prepare the single cell libraries. First, during the process of tagmentation the cDNAs are fragmented and tagged with adapter sequences by a transposase. Then, the tagmented cDNAs were amplified using a unique combination of i5 and i7 indexes and purified using AMPure XP beads (Beckman Coulter). To control for the appropriate size distribution of 250-1000bp of the libraries a high sensitivity bioanalyzer (Agilent Technologies) was used and the concentration again determined via Qubit. The libraries were pooled at a concentration of 1nM, denatured with 0.2 N NaOH and diluted to 20pM. The sequencing was run on a NextSeq 550 (Illumina).

### scRNAseq analysis

#### Sequencing quality control, mapping and expression quantification

Sequences were demultiplexed using bcl2fastq v2.20.0 and trimmed to remove adapters and low quality end bases using cutadapt v2.9. After trimming, reads shorter than 20 bases were discarded. Reads were then aligned to the *mm10* mouse reference genome using STAR (v2.7.9a) with default options (Dobin et al., 2013). Mapped reads were then quantitated using the RSEM pipeline (v1.3.3) with the *--single-cell-prior* option (2).

#### Data filtering

Quality control and data filtering were performed separately for the two sequencing runs. In both runs, cells with more than 10 million reads or 20% of reads mapping to mitonchondrial genes or more than 1% of spike-ins reads were filtered out. Cells expressing fewer than 4800 genes in the sequencing run 1 or fewer than 4,000 genes in the sequencing run 2 were excluded from the downstream analyses. 170 cells out of the 184 sequenced cells (20× E3.25, 22× E3.75, 14× E4.0, 31× E2.75+16h ctl, 31× E2.75+16h LY, 28× E2.75+24h ctl, 24× E2.75+24h LY) were kept for downstream analyses. Reads were aligned to the mm10 genome and transcripts per million (TPM) computed. TPM counts for each sample are available in Supplemental Figure 7.

#### Data integration and downstream analyses

Sequenced cells were integrated with data from previously published datasets (Mohammed et al., 2017; Nowotschin et al., 2019; Posfai et al., 2017). Before integration, a check for quality control and a data filtering step was performed on each individual run from Nowotschin et al. (Nowotschin et al., 2019). The different thresholds used for each of these runs are detailed in Supplemental Table 6. The datasets were normalized as counts per million, log-transformed and then integrated using Scanorama version 1.7.1 (Hie et al., 2019). Variable genes in each individual datasets were identified by computing the standardized variance for each gene and the 158 genes tagged as variable in at least 4 different sequencing runs were used for data integration. Cells were then clustered using the standard workflow from Seurat version 4.1.0 (Butler et al., 2018). Differentially expressed genes were identified using the MAST version 1.22.0 (Finak et al., 2015) accounting for the data origin. Using the AddModuleScore function from Seurat with default parameters, the average mean expressions of the 20 genes with the highest fold changes and the 20 genes with the lowest fold changes when comparing vehicle and LY-treated cells were scored in all cells from the external datasets clustered in the ICM.

---

**Figure S1:**
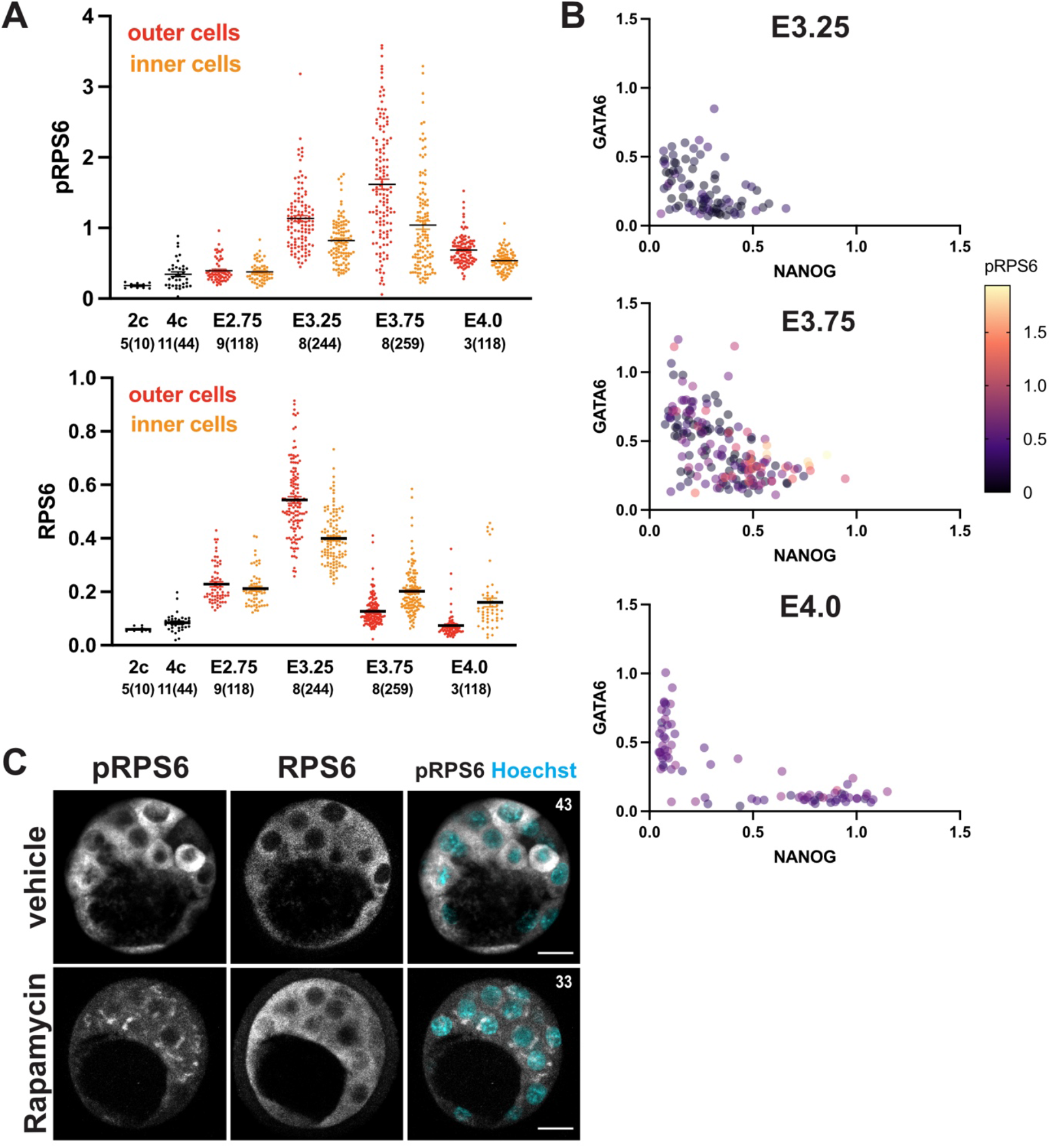
pRPS6 levels show no correlation with NANOG and GATA6 during blastocyst stages. (A) Quantification of pRPS6 (left) and RPS6 (right) levels in inner and outer cells from 2 cell stage to E4.0. Number of embryos and analyzed cells per stage indicated at the bottom. (B) Level of nuclear GATA6/NANOG and cytoplasmic pRPS6 in inner cells of E3.25, E3.75 and E4.0 embryos. (C) Immunodetection of pRPS6 and RPS6 (grey) in embryos cultured from E2.75 for 24h in mTORC1 inhibitor Rapamycin. Data are from one experiment, representative of two independent experiments. Pictures do correspond to the projection of 5 optical slices. Number of cells per embryo indicated on the top right of images. Scale bar: 20µm.

**Figure S2:**
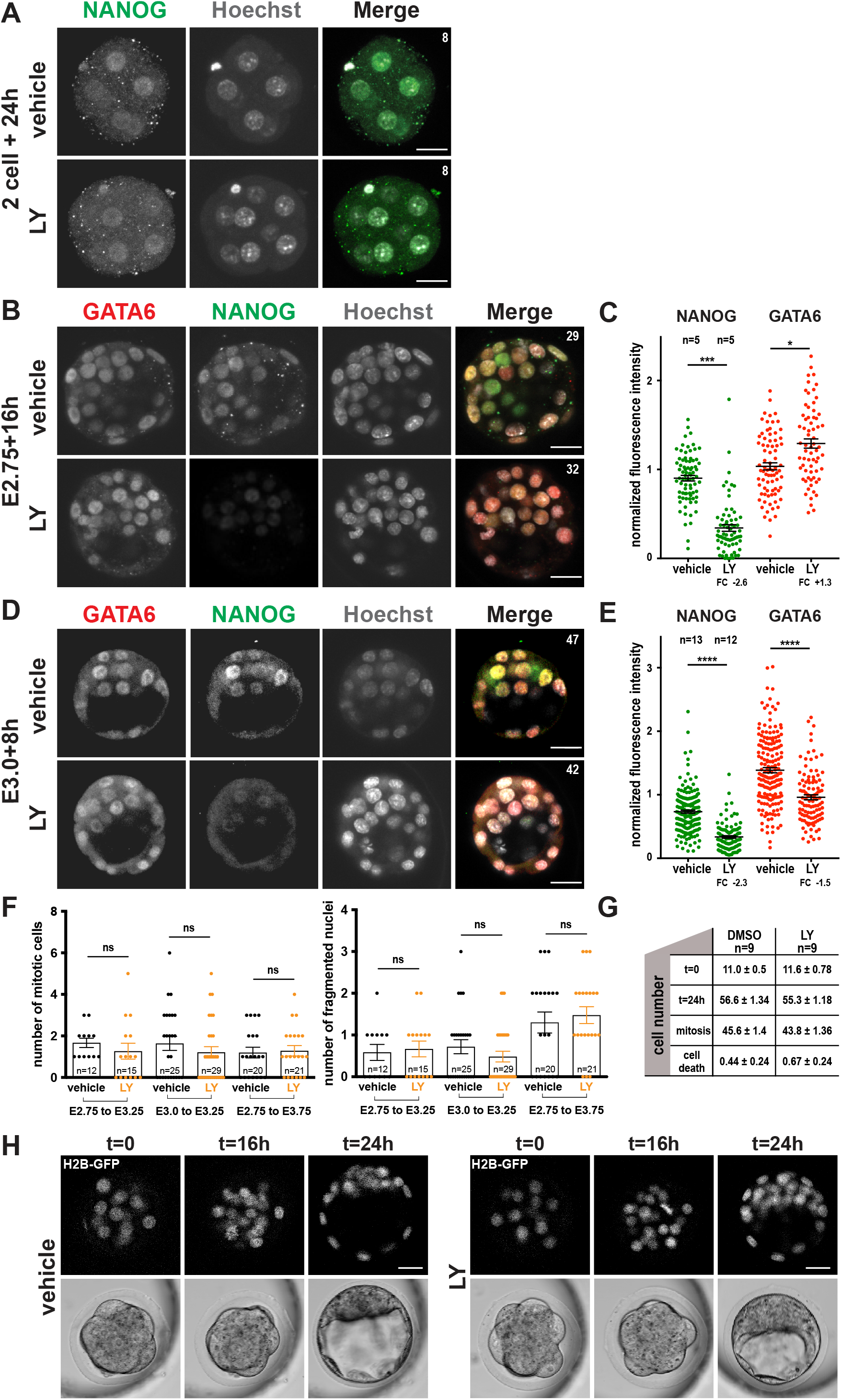
PI3K/AKT regulates NANOG through survival and proliferation independent mechanisms. (A) Immunodetection of NANOG (green) in embryos cultured for 24h from the 2-cell stage. (B) Immunodetection and (C) quantification of GATA6 (red) and NANOG (green) in embryos cultured for 16h from E2.75. (D) Immunodetection and (E) quantification of GATA6 (red) and NANOG (green) in embryos cultured for 8h from E3.0. (F) Number of mitotic cells (left) and cells undergoing cell death (right) in fixed embryos cultured for different durations in absence or presence of LY. Graphs show the results of three independent experiments. (G) Total cell number at the start and the end of live imaging, cumulative number of mitotic cells and of cells undergoing cell death and (H) images of vehicle and LY treated embryos expressing H2B-GFP at the start of live imaging (t=0), after 16h (t=16h) and after 24h (t=24h). Data are from one experiment, representative of at least two (A, H) and more than 4 (B-E) independent experiments. Pictures do correspond to the projection of 10 (A) or 5 (B, D, H) optical slices. Number of cells per embryo indicated on the top right of images. N: number of embryos.

**Figure S3:**
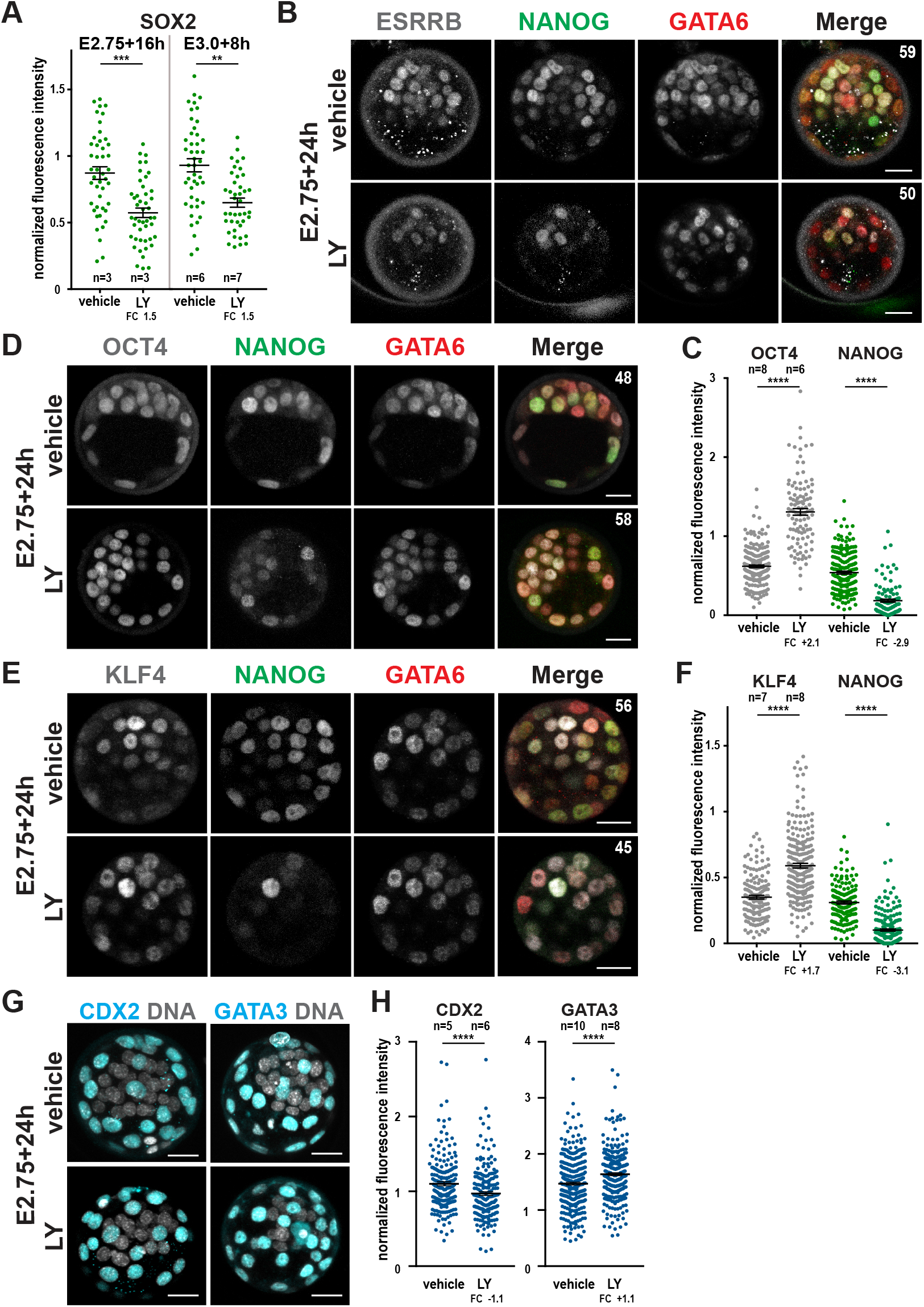
No reduction of Epi markers OCT4 and KLF4 and the TE markers CDX2 and GATA3 upon PI3K inhibition. (A) Quantification of SOX2 (green) in embryos cultured for 16h from E2.75 or 8h from E3.0. (B) Immunodetection of ESRRB (grey), GATA6 (red) and NANOG (green) in embryos cultured for 24h from E2.75. (C, E) Immunodetection and (D, F) quantification of OCT4 (grey), KLF4 (grey), GATA6 (red) and NANOG (green) in embryos cultured for 24h from E2.75. (G, H) Immunodetection and quantification of CDX2 (right) and GATA3 (left) in TE cells of embryos cultured for 24h from E2.75. Data are from one experiment, representative of two (C-F) or more than three (A, G-H) independent experiments. Pictures do correspond to the projection of 5 optical slices. Number of cells per embryo indicated on the top right of images. Scale bar: 20µm; n: number of embryos; FC: fold change.

**Figure S4:**
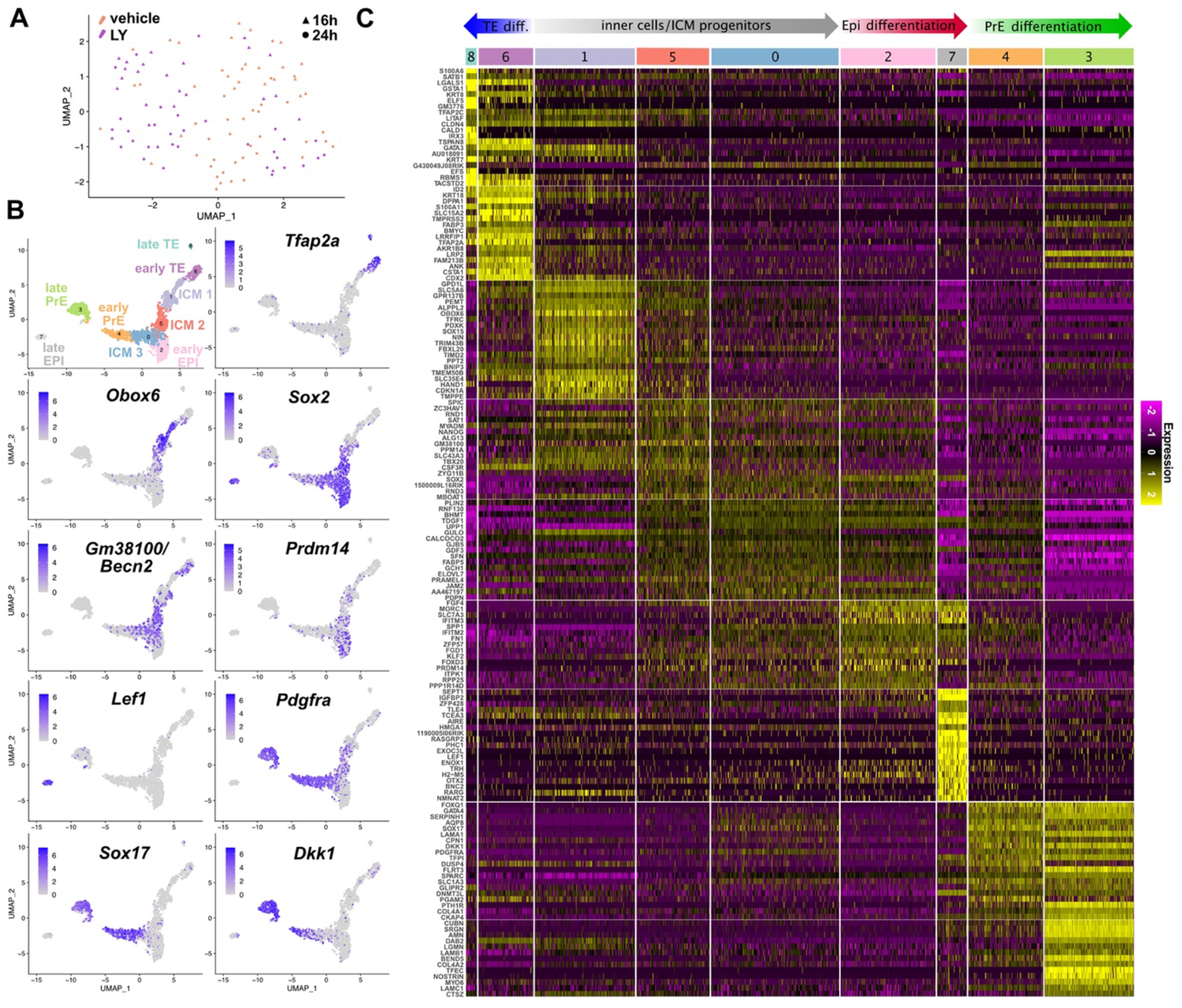
Single cell transcriptome profiling of morula to late blastocyst mouse embryos. (A) UMAP of vehicle and LY-treated cells. (B) UMAPs of all cells colored by gene expression level of markers enriched in the different clusters. Grey to purple scale indicates average expression. (C) Heatmap of differentially expressed genes between the 9 identified clusters. Top 20 enriched markers ranked according to the p value are indicated for each cluster.

**Figure S5:**
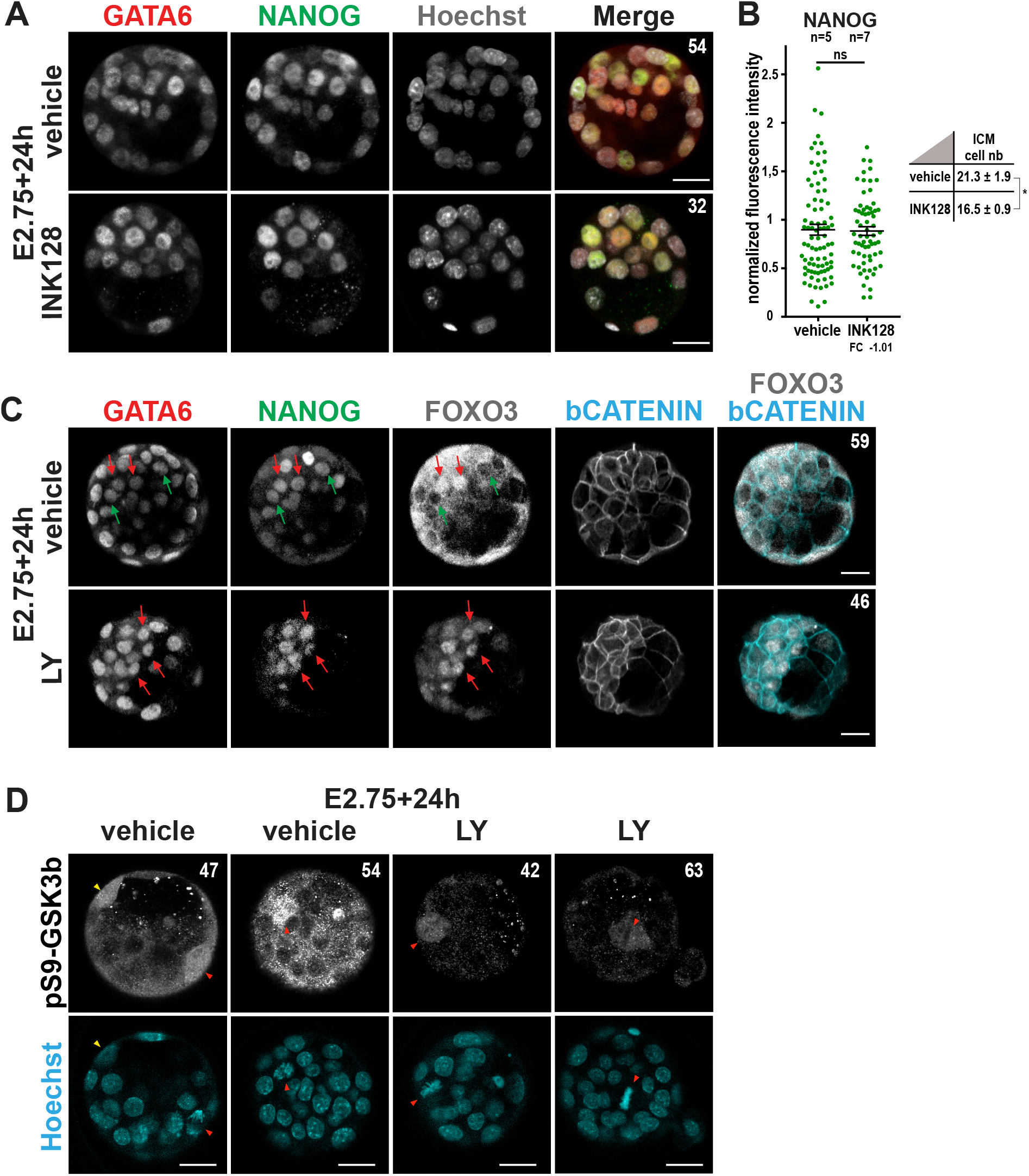
NANOG levels are not regulated by mTOR and FOXO3. (A) Immunodetection of GATA6 (red) and NANOG (green) and (B) quantification of NANOG (green) in embryos cultured for 24h from E2.75. Embryos cultured with INK128 showed reduced cell number with no sign of cell death probably due to its known pausing action on cellular metabolism (Bulut-Karslioglu et al., 2016; Xu et al., 2021). (C) Immunodetection of GATA6 (red), NANOG (green), FOXO3 (grey) and β-CATENIN (cyan) in embryos cultured for 24h from E2.75. Green arrows point to examples of inner cells with cytoplasmic FOXO3. Red arrows point to examples of inner cells with nuclear FOXO3. Data are from one experiment, representative of three independent experiments. Pictures do correspond to the projection of 5 optical slices. (D) Immuodetection of pS9-GSK3b (gray) in embryos cultured for 24h from E2.75. Pictures do correspond a single optical slice. Red arrowheads point to mitotic cells located in the z stacks shown while the yellow arrowhead points to a mitotic outer cell extending over several satcks. Number of cells per embryo indicated on the top right of images. Scale bar: 20µm; n: number of embryos; FC: fold change.

**Figure S6:**
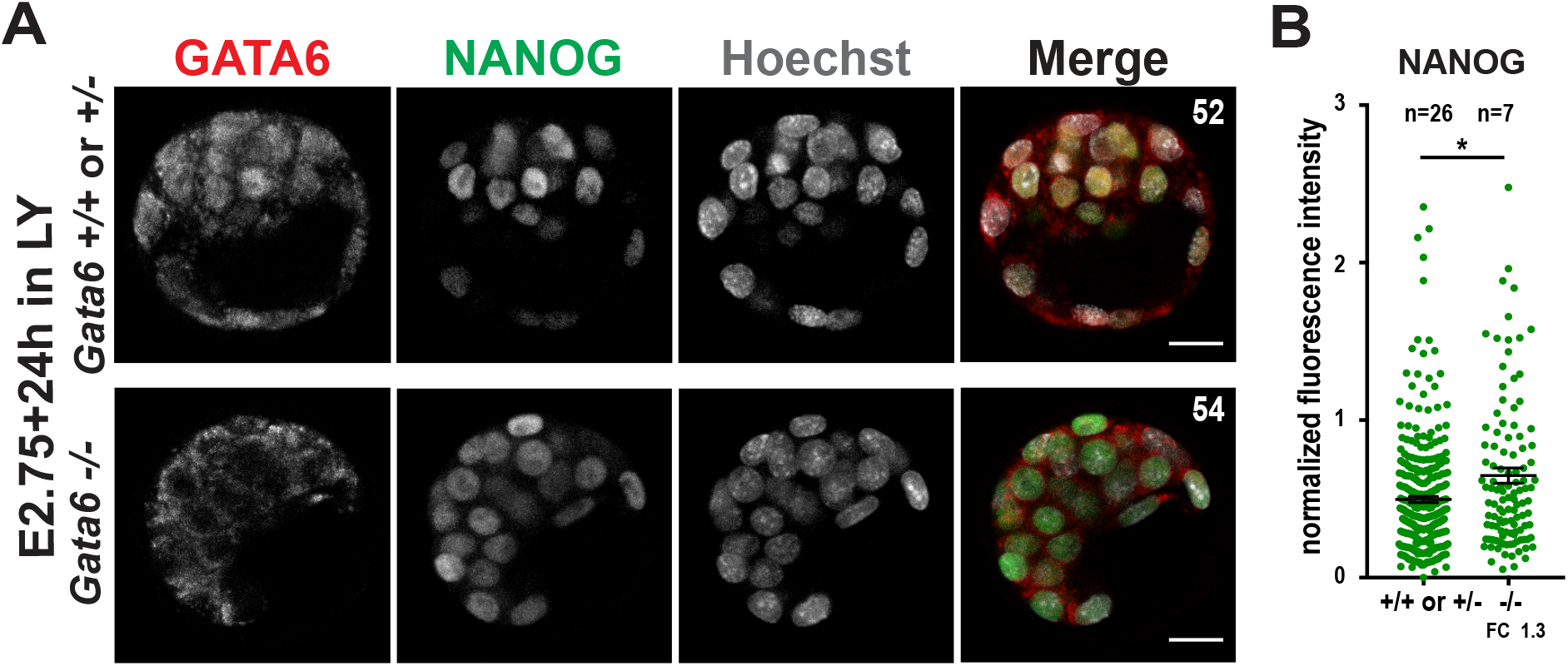
Effect of PI3K inhibition on NANOG levels in zygotic *Gata6* mutant embryos. (A) Immunodetection of GATA6 (red) of NANOG (green) and (B) quantification of NANOG (green) in zygotic *Gata6 +/+, +/- and -/-* embryos cultured for 24h from E2.75. Graphs shows results from three independent experiments. Pictures do correspond to the projection of 5 optical slices. Number of cells per embryo indicated on the top right of images. Scale bar: 20µm; n: number of embryos; FC: fold change.

## Supplemental Tables

**Table S1.** List of genes enriched in each cluster of the UMAP of all integrated cells over all other clusters.

**Table S2.** List of DEG in ICM cells from vehicle versus LY-treated embryos.

**Table S3.** List of compounds used in this study.

**Table S4.** List of antibodies used in this study.

**Table S5.** List of oligonucleotides used in this study.

**Table S6.** Thresholds used for the different runs from Nowotschin et al, 2019.

## Notes

### Competing Interest Statement

The authors have declared no competing interest.

## REFERENCES

Aguirre-Lavin, T., Adenot, P., Bonnet-Garnier, A., Lehmann, G., Fleurot, R., Boulesteix, C., Debey, P. and Beaujean, N. (2012). 3D-FISH analysis of embryonic nuclei in mouse highlights several abrupt changes of nuclear organization during preimplantation development. Bmc Dev Biol 12, 30.

Allègre, N., Chauveau, S., Dennis, C., Renaud, Y., Meistermann, D., Estrella, L. V., Pouchin, P., Cohen-Tannoudji, M., David, L. and Chazaud, C. (2022). NANOG initiates epiblast fate through the coordination of pluripotency genes expression. Nat Commun 13, 3550.

Artus, J., Piliszek, A. and Hadjantonakis, A.-K. K. (2011). The primitive endoderm lineage of the mouse blastocyst: sequential transcription factor activation and regulation of differentiation by Sox17. Developmental Biology 350, 393–404.

Azami, T., Bassalert, C., Allègre, N., Estrella, L. V., Pouchin, P., Ema, M. and Chazaud, C. (2019). Regulation of the ERK signalling pathway in the developing mouse blastocyst. Development 146, dev177139.

Bechard, M. and Dalton, S. (2009). Subcellular localization of glycogen synthase kinase 3beta controls embryonic stem cell self-renewal. Mol Cell Biol 29, 2092–104.

Bessonnard, S., Mot, L. D., Gonze, D., Barriol, M., Dennis, C., Goldbeter, A., Dupont, G. and Chazaud, C. (2014). Gata6, Nanog and Erk signaling control cell fate in the inner cell mass through a tristable regulatory network. Development 141, 3637–3648.

Bessonnard, S., Coqueran, S., Vandormael-Pournin, S., Dufour, A., Artus, J. and Cohen-Tannoudji, M. (2017). ICM conversion to epiblast by FGF/ERK inhibition is limited in time and requires transcription and protein degradation. Sci. Rep. 7, 12285.

Bessonnard, S., Vandormael-Pournin, S., Coqueran, S., Cohen-Tannoudji, M. and Artus, J. (2019). PDGF Signaling in Primitive Endoderm Cell Survival Is Mediated by PI3K-mTOR Through p53-Independent Mechanism. Stem Cells 37, 888–898.

Borensztein, M., Syx, L., Servant, N. and Heard, E. (2018). Transcriptome Profiling of Single Mouse Oocytes. Methods Mol. Biol. 1818, 51–65.

Boroviak, T., Loos, R., Lombard, P., Okahara, J., Behr, R., Sasaki, E., Nichols, J., Smith, A. G. and Bertone, P. (2015). Lineage-Specific Profiling Delineates the Emergence and Progression of Naive Pluripotency in Mammalian Embryogenesis. Dev. Cell 35, 366– 382.

Brunet, A., Bonni, A., Zigmond, M. J., Lin, M. Z., Juo, P., Hu, L. S., Anderson, M. J., Arden, K. C., Blenis, J. and Greenberg, M. E. (1999). Akt Promotes Cell Survival by Phosphorylating and Inhibiting a Forkhead Transcription Factor. Cell 96, 857–868.

Bulut-Karslioglu, A., Biechele, S., Jin, H., Macrae, T. A., Hejna, M., Gertsenstein, M., Song, J. S. and Ramalho-Santos, M. (2016). Inhibition of mTOR induces a paused pluripotent state. Nature 540, 119–123.

Butler, A., Hoffman, P., Smibert, P., Papalexi, E. and Satija, R. (2018). Integrating single-cell transcriptomic data across different conditions, technologies, and species. Nat Biotechnol 36, 411–420.

Campbell, J. M., Nottle, M. B., Vassiliev, I., Mitchell, M. and Lane, M. (2012). Insulin increases epiblast cell number of in vitro cultured mouse embryos via the PI3K/GSK3/p53 pathway. Stem Cells and Development 21, 2430–2441.

Cartwright, P., McLean, C., Sheppard, A., Rivett, D., Jones, K. and Dalton, S. (2005). LIF/STAT3 controls ES cell self-renewal and pluripotency by a Myc-dependent mechanism. Development 132, 885–896.

Chazaud, C., Yamanaka, Y., Pawson, T. and Rossant, J. (2006). Early lineage segregation between epiblast and primitive endoderm in mouse blastocysts through the Grb2-MAPK pathway. Dev. Cell 10, 615–624.

Chen, B., Xue, Z., Yang, G., Shi, B., Yang, B., Yan, Y., Wang, X., Han, D., Huang, Y. and Dong, W. (2013). Akt-Signal Integration Is Involved in the Differentiation of Embryonal Carcinoma Cells. Plos One 8, e64877.

Cross, D. A. E., Alessi, D. R., Cohen, P., Andjelkovich, M. and Hemmings, B. A. (1995). Inhibition of glycogen synthase kinase-3 by insulin mediated by protein kinase B. Nature 378, 785–789.

Deng, Q., Ramsköld, D., Reinius, B. and Sandberg, R. (2014). Single-cell RNA-seq reveals dynamic, random monoallelic gene expression in mammalian cells. Science 343, 193– 196.

Dobin, A., Davis, C. A., Schlesinger, F., Drenkow, J., Zaleski, C., Jha, S., Batut, P., Chaisson, M. and Gingeras, T. R. (2013). STAR: ultrafast universal RNA-seq aligner. Bioinformatics 29, 15–21.

Fang, L., Zhang, L., Wei, W., Jin, X., Wang, P., Tong, Y., Li, J., Du, J. X. and Wong, J. (2014). A methylation-phosphorylation switch determines Sox2 stability and function in ESC maintenance or differentiation. Molecular Cell 55, 537–551.

Festuccia, N., Vandormael-Pournin, S., Chervova, A., Geiselman, A., Langa-Vives, F., Coux, R.-X., Gonzalez, I., Cohen-Tannoudji, M. and Navarro, P. (2023). Nr5a2 is essential for morula development. Biorxiv 2023.01.16.524255.

Finak, G., McDavid, A., Yajima, M., Deng, J., Gersuk, V., Shalek, A. K., Slichter, C. K., Miller, H. W., McElrath, M. J., Prlic, M., et al. (2015). MAST: a flexible statistical framework for assessing transcriptional changes and characterizing heterogeneity in single-cell RNA sequencing data. Genome Biol 16, 278.

Frankenberg, S., Gerbe, F., Bessonnard, S., Belville, C., Pouchin, P., Bardot, O. and Chazaud, C. (2011). Primitive endoderm differentiates via a three-step mechanism involving Nanog and RTK signaling. Dev. Cell 21, 1005–1013.

Goolam, M., Scialdone, A., Graham, S. J. L., Macaulay, I. C., Jedrusik, A., Hupalowska, A., Voet, T., Marioni, J. C. and Zernicka-Goetz, M. (2016). Heterogeneity in Oct4 and Sox2 Targets Biases Cell Fate in 4-Cell Mouse Embryos. Cell 165, 61–74.

Guo, G., Huss, M., Tong, G. Q., Wang, C., Sun, L. L., Clarke, N. D. and Robson, P. (2010). Resolution of Cell Fate Decisions Revealed by Single-Cell Gene Expression Analysis from Zygote to Blastocyst. Dev Cell 18, 675–685.

Hadjantonakis, A.-K. and Papaioannou, V. E. (2004). Dynamic in vivo imaging and cell tracking using a histone fluorescent protein fusion in mice. Bmc Biotechnol 4, 33.

Hahaut, V., Pavlinic, D., Carbone, W., Schuierer, S., Balmer, P., Quinodoz, M., Renner, M., Roma, G., Cowan, C. S. and Picelli, S. (2022). Fast and highly sensitive full-length single-cell RNA sequencing using FLASH-seq. Nat Biotechnol 40, 1447–1451.

Halet, G., Viard, P. and Carroll, J. (2008). Constitutive PtdIns(3,4,5)P3 synthesis promotes the development and survival of early mammalian embryos. Development 135, 425– 429.

Hamazaki, T., Kehoe, S. M., Nakano, T. and Terada, N. (2006). The Grb2/Mek pathway represses Nanog in murine embryonic stem cells. Mol. Cell. Biol. 26, 7539–7549.

Hie, B., Bryson, B. and Berger, B. (2019). Efficient integration of heterogeneous single-cell transcriptomes using Scanorama. Nat Biotechnol 37, 685–691.

Hishida, T., Nakachi, Y., Mizuno, Y., Katano, M., Okazaki, Y., Ema, M., Takahashi, S., Hirasaki, M., Suzuki, A., Ueda, A., et al. (2015). Functional Compensation Between Myc and PI3K Signaling Supports Self-Renewal of Embryonic Stem Cells. Stem Cells 33, 713–725.

Kang, M., Piliszek, A., Artus, J. and Hadjantonakis, A.-K. K. (2013). FGF4 is required for lineage restriction and salt-and-pepper distribution of primitive endoderm factors but not their initial expression in the mouse. Development 140, 267–279.

Kang, M., Garg, V. and Hadjantonakis, A.-K. K. (2017). Lineage Establishment and Progression within the Inner Cell Mass of the Mouse Blastocyst Requires FGFR1 and FGFR2. Dev. Cell 41, 496–510.e5.

Lallemand, Y., Luria, V., Haffner-Krausz, R. and Lonai, P. (1998). Maternally expressed PGK-Cre transgene as a tool for early and uniform activation of the Cre site-specific recombinase. Transgenic Res 7, 105–112.

Lin, T.-C., Yen, J.-M., Gong, K.-B., Hsu, T.-T. and Chen, L.-R. (2003). IGF-1/IGFBP-1 increases blastocyst formation and total blastocyst cell number in mouse embryo culture and facilitates the establishment of a stem-cell line. BMC Cell Biol. 4, 14.

Lin, Y., Yang, Y., Li, W., Chen, Q., Li, J., Pan, X., Zhou, L., Liu, C., Chen, C., He, J., et al. (2012). Reciprocal regulation of Akt and Oct4 promotes the self-renewal and survival of embryonal carcinoma cells. Molecular Cell 48, 627–640.

Lu, D. P., Chandrakanthan, V., Cahana, A., Ishii, S. and O’Neill, C. (2004). Trophic signals acting via phosphatidylinositol-3 kinase are required for normal pre-implantation mouse embryo development. J Cell Sci 117, 1567–1576.

Mendoza, M. C., Er, E. E. and Blenis, J. (2011). The Ras-ERK and PI3K-mTOR pathways: cross-talk and compensation. Trends in Biochemical Sciences 36, 320–328.

Meyuhas, O. (2015). Chapter Two Ribosomal Protein S6 Phosphorylation Four Decades of Research.pp. 41–73. International Review of Cell and Molecular Biology.

Mohammed, H., Hernando-Herraez, I., Savino, A., Scialdone, A., Macaulay, I., Mulas, C., Chandra, T., Voet, T., Dean, W., Nichols, J., et al. (2017). Single-Cell Landscape of Transcriptional Heterogeneity and Cell Fate Decisions during Mouse Early Gastrulation. Cell Reports 20, 1215–1228.

Molotkov, A., Mazot, P., Brewer, J. R., Cinalli, R. M. and Soriano, P. (2017). Distinct Requirements for FGFR1 and FGFR2 in Primitive Endoderm Development and Exit from Pluripotency. Dev. Cell 41, 511–526.e4.

Morgani, S. M. and Brickman, J. M. (2015). LIF supports primitive endoderm expansion during pre-implantation development. Development 142, 3488–3499.

Mot, L. D., Gonze, D., Bessonnard, S., Chazaud, C., Goldbeter, A. and Dupont, G. (2016). Cell Fate Specification Based on Tristability in the Inner Cell Mass of Mouse Blastocysts. Biophysical Journal 110, 710–722.

Nichols, J., Silva, J. C. R., Roode, M. and Smith, A. G. (2009). Suppression of Erk signalling promotes ground state pluripotency in the mouse embryo. Development 136, 3215– 3222.

Niwa, H., Ogawa, K., Shimosato, D. and Adachi, K. (2009). A parallel circuit of LIF signalling pathways maintains pluripotency of mouse ES cells. Nature 460, 118–122.

Nowotschin, S., Setty, M., Kuo, Y.-Y., Liu, V., Garg, V., Sharma, R., Simon, C. S., Saiz, N., Gardner, R., Boutet, S. C., et al. (2019). The emergent landscape of the mouse gut endoderm at single-cell resolution. Nature 569, 361–367.

Okamura, E., Tam, O. H., Posfai, E., Li, L., Cockburn, K., Lee, C. Q. E., Garner, J. and Rossant, J. (2019). Esrrb function is required for proper primordial germ cell development in presomite stage mouse embryos. Dev Biol 455, 382–392.

Paling, N. R. D., Wheadon, H., Bone, H. K. and Welham, M. J. (2004). Regulation of embryonic stem cell self-renewal by phosphoinositide 3-kinase-dependent signaling. J. Biol. Chem. 279, 48063–48070.

Plusa, B., Piliszek, A., Frankenberg, S., Artus, J. and Hadjantonakis, A.-K. K. (2008). Distinct sequential cell behaviours direct primitive endoderm formation in the mouse blastocyst. Development 135, 3081–3091.

Posfai, E., Petropoulos, S., Barros, F. R. O. de, Schell, J. P., Jurisica, I., Sandberg, R., Lanner, F. and Rossant, J. (2017). Position- and Hippo signaling-dependent plasticity during lineage segregation in the early mouse embryo. Elife 6, e22906.

Ramos-Ibeas, P., Sang, F., Zhu, Q., Tang, W. W. C., Withey, S., Klisch, D., Wood, L., Loose, M., Surani, M. A. and Alberio, R. (2019). Pluripotency and X chromosome dynamics revealed in pig pre-gastrulating embryos by single cell analysis. Nat Commun 10, 500.

Riley, J. K., Carayannopoulos, M. O., Wyman, A. H., Chi, M., Ratajczak, C. K. and Moley, K. H. (2005). The PI3K/Akt pathway is present and functional in the preimplantation mouse embryo. Developmental Biology 284, 377–386.

Saiz, N., Williams, K. M., Seshan, V. E. and Hadjantonakis, A.-K. K. (2016). Asynchronous fate decisions by single cells collectively ensure consistent lineage composition in the mouse blastocyst. Nature Communications 7, 13463.

Saiz, N., Mora-Bitria, L., Rahman, S., George, H., Herder, J. P., Garcia-Ojalvo, J. and Hadjantonakis, A.-K. (2019). Growth-factor-mediated coupling between lineage size and cell fate choice underlies robustness of mammalian development. Elife 9, e56079.

Sanchez-Ripoll, Y., Bone, H. K., Owen, T., Guedes, A. M. V., Abranches, E., Kumpfmueller, B., Spriggs, R. V., Henrique, D. and Welham, M. J. (2013). Glycogen Synthase Kinase-3 Inhibition Enhances Translation of Pluripotency-Associated Transcription Factors to Contribute to Maintenance of Mouse Embryonic Stem Cell Self-Renewal. Plos One 8, e60148.

Santostefano, K. E., Hamazaki, T., Pardo, C. E., Kladde, M. P. and Terada, N. (2012). Fibroblast growth factor receptor 2 homodimerization rapidly reduces transcription of the pluripotency gene Nanog without dissociation of activating transcription factors. Journal of Biological Chemistry 287, 30507–30517.

Schrode, N., Saiz, N., Talia, S. D. and Hadjantonakis, A.-K. K. (2014). GATA6 levels modulate primitive endoderm cell fate choice and timing in the mouse blastocyst. Dev. Cell 29, 454–467.

Singh, A. M., Bechard, M., Smith, K. and Dalton, S. (2012). Reconciling the different roles of Gsk3β in “naïve” and “primed” pluripotent stem cells. Cell Cycle 11, 2991–2996.

Smith, K. N., Singh, A. M. and Dalton, S. (2010). Myc Represses Primitive Endoderm Differentiation in Pluripotent Stem Cells. Cell Stem Cell 7, 343–354.

Sodhi, C. P., Li, J. and Duncan, S. A. (2006). Generation of mice harbouring a conditional loss-of-function allele of Gata6. Bmc Dev Biol 6, 19–19.

Storm, M. P., Bone, H. K., Beck, C. G., Bourillot, P.-Y., Schreiber, V., Damiano, T., Nelson, A., Savatier, P. and Welham, M. J. (2007). Regulation of Nanog expression by phosphoinositide 3-kinase-dependent signaling in murine embryonic stem cells. J. Biol. Chem. 282, 6265–6273.

Storm, M. P., Kumpfmueller, B., Thompson, B., Kolde, R., Vilo, J., Hummel, O., Schulz, H. and Welham, M. J. (2009). Characterization of the Phosphoinositide 3-Kinase-Dependent Transcriptome in Murine Embryonic Stem Cells: Identification of Novel Regulators of Pluripotency. Stem Cells 27, 764–775.

Tang, F., Barbacioru, C., Nordman, E., Li, B., Xu, N., Bashkirov, V. I., Lao, K. and Surani, M. A. (2010). RNA-Seq analysis to capture the transcriptome landscape of a single cell. Nature Protocols 5, 516–535.

Tosenberger, A., Gonze, D., Bessonnard, S., Cohen-Tannoudji, M., Chazaud, C. and Dupont, G. (2017). A multiscale model of early cell lineage specification including cell division. Npj Syst Biology Appl 3, 16.

Vandormael-Pournin, S., Frachon, E., Gobaa, S. and Cohen-Tannoudji, M. (2021). Microfabricated Device for High-Resolution Imaging of Preimplantation Embryos. Methods Mol. Biol. 2214, 11–30.

Vries, W. N. de, Binns, L. T., Fancher, K. S., Dean, J., Moore, R., Kemler, R. and Knowles, B. B. (2000). Expression of Cre recombinase in mouse oocytes: a means to study maternal effect genes. Genesis 26, 110–112.

Wamaitha, S. E., Grybel, K. J., Alanis-Lobato, G., Gerri, C., Ogushi, S., McCarthy, A., Mahadevaiah, S. K., Healy, L., Lea, R. A., Molina-Arcas, M., et al. (2020). IGF1-mediated human embryonic stem cell self-renewal recapitulates the embryonic niche. Nat Commun 11, 764.

Wicklow, E., Blij, S., Frum, T., Hirate, Y., Lang, R. A., Sasaki, H. and Ralston, A. (2014). HIPPO Pathway Members Restrict SOX2 to the Inner Cell Mass Where It Promotes ICM Fates in the Mouse Blastocyst. Plos Genet 10, e1004618.

Wray, J., Kalkan, T., Gomez-Lopez, S., Eckardt, D., Cook, A., Kemler, R. and Smith, A. (2011). Inhibition of glycogen synthase kinase-3 alleviates Tcf3 repression of the pluripotency network and increases embryonic stem cell resistance to differentiation. Nat Cell Biol 13, 838–845.

Xu, X., Ahmed, T., Wang, L., Cao, X., Zhang, Z., Wang, M., Lv, Y., Kanwal, S., Tariq, M., Lin, R., et al. (2021). The mTORC1-eIF4F axis controls paused pluripotency. EMBO reports e53081.

Yamanaka, Y., Lanner, F. and Rossant, J. (2010). FGF signal-dependent segregation of primitive endoderm and epiblast in the mouse blastocyst. Development 137, 715–724.

Yi, F., Pereira, L., Hoffman, J. A., Shy, B. R., Yuen, C. M., Liu, D. R. and Merrill, B. J. (2011). Opposing effects of Tcf3 and Tcf1 control Wnt stimulation of embryonic stem cell self-renewal. Nature Cell Biology 13, 762–770.

Yu, J. S. L. and Cui, W. (2016). Proliferation, survival and metabolism: the role of PI3K/AKT/mTOR signalling in pluripotency and cell fate determination. Development 143, 3050–3060.

Zhang, X., Yalcin, S., Lee, D.-F., Yeh, T.-Y. J., Lee, S.-M., Su, J., Mungamuri, S. K., Rimmelé, P., Kennedy, M., Sellers, R., et al. (2011). FOXO1 is an essential regulator of pluripotency in human embryonic stem cells. Nature Cell Biology 13, 1092–1099.

Zimmermann, S. and Moelling, K. (1999). Phosphorylation and regulation of Raf by Akt (protein kinase B). Science 286, 1741–1744.

